# A neurofunctional signature of romantic love predicts rewards on social media, effects of oxytocin and drug-cue reactivity

**DOI:** 10.64898/2026.08.10.744060

**Authors:** Yu Wu, Xianyang Gan, Guojuan Jiao, Rene Hurlemann, Dirk Scheele, Xinqi Zhou, Ran Zhang, Feng Zhou, Heng Jiang, Kun Fu, Junjie Wang, Yue Teng, Benjamin Becker

**Author notes:** Corresponding author: Benjamin Becker, Department of Psychology, The University of Hong Kong, Hong Kong SAR, 999077, China.

## Abstract

Romantic love is a selective motivational state supporting pair bonding, yet its neural representation and relation to other affiliative-rewarding experiences remain unclear. Across eight fMRI studies (*n*=420) spanning naturalistic affiliative-reward, pharmacological and addiction-relevant experiments, we developed and evaluated multivariate whole-brain decoders of romantic love and friendship. Both signatures engaged mesocorticolimbic reward systems and the precuneus, yet were dissociable at the whole-brain level and in their recruitment of social-cognitive systems. The love signature generalized to reward on social media, was specific to positive valence, and did not track sweet-taste or monetary reward, indicating a distinct social-affiliative reward representation. Oxytocin selectively increased love - but not friendship - signature reactivity to the romantic partner. In an independent drug-cue-reactivity dataset, the love signature specifically identified heavy cannabis users. These findings establish a dissociable neurofunctional signature of romantic love in conserved bonding circuits that extend to digital affiliation and are co-opted in addiction.

## Introduction

Pair-bonding - the capacity to form an enduring, selective bond with a single partner - is evolutionarily conserved across socially monogamous species, from prairie voles to humans, and likely arose through specialization of the systems mediating broader affiliative attachments^1,2^. In humans, this bond is experienced as romantic love and is among the most powerful and selective of motivational drives, whereas other affiliative bonds such as friendship subserve broader functions like non-kin cooperation^1^. Rodent models indicate that this selective motivational quality depends on integrated dopaminergic and oxytocinergic signaling within mesocorticolimbic reward circuitry^3,4^ - a neural basis that, together with behavioral similarities such as preoccupation, craving, and resistance to substitution, has motivated hypotheses linking romantic love to addiction^5,6^ and, more recently, to engagement with social media and AI^7,8^. Yet a comprehensive and precise neurofunctional signature of romantic love in humans remains elusive, impeding empirical tests of these theories. Here, we develop an fMRI-based whole-brain signature of romantic love, distinguish it from friendship, and show that it is selectively sensitive to neural representations of substance addiction and oxytocin and generalizes to social-media reward, yet remains distinct from nonspecific non-social rewards (sweet taste and monetary reward).

Insights into the neurobiology of highly selective pair bonds have been driven by discoveries in socially monogamous prairie voles, which form long-lasting bonds. This work shows that selective pair-bond formation depends on interactions between oxytocinergic, vasopressinergic and mesocorticolimbic dopamine systems, with the nucleus accumbens and medial prefrontal cortex playing central roles in linking partner-specific social representations to reward^2,9–11^. Together with findings in other model organisms, these data suggest that pair-bonding - and, in humans, romantic love - is built upon an evolutionarily conserved mesocorticolimbic circuitry that also mediates more ancient forms of social affiliation as well as general reward and reinforcement, yet nonetheless possesses a partly distinguishable neural signature^1,12^. In humans, the neural basis of romantic love has been examined primarily with functional magnetic resonance imaging (fMRI), as individuals view or anticipate the face of their romantic partner relative to a familiar other. Romantic-partner cues reliably engage core nodes of the mesocorticolimbic circuitry - including the nucleus accumbens and medial frontal regions - across cultures and relationship durations^13–17^, converging on a conserved reward circuitry that may constitute a core substrate of romantic love (see also refs. 1 and 2).

However, these exposure paradigms capture only a narrow, stimulus-driven facet of romantic love, which extends beyond the immediate presence of the partner to the relationship-defining positive memories that sustain the bond^18^. Moreover, ventral striatum and medial prefrontal responses are observed across non-romantic bonds, including mother-infant bonds^19^ and friendships or other close relationships^20–22^, indicating that these regions are not specific to romantic attachment. Recent meta-analyses further report that affiliative stimuli engage an extensive common architecture encompassing the ventral striatum, medial frontal cortex, and posterior medial cortex^23^ - a network that overlaps with regions processing non-social reinforcers such as money^13,24,25^ - while experimental work employing direct comparisons among different affiliative bonds indicates engagement of partly dissociable neural systems^26,27^. It therefore remains unclear whether the neural representation of romantic love can be distinguished from that of other social-affiliative or non-social rewards in humans.

While love and enduring relationships confer well-documented benefits, the same reward and motivational circuits that sustain them can also drive maladaptive, excessive, and ultimately compulsive behavior, including problematic social-media use and escalating drug use. Accumulating evidence shows that online social interactions, such as receiving “likes” and positive feedback, engage the ventral striatum and associated medial prefrontal regions^28–31^. These regions overlap with mesocorticolimbic nodes that, in animal models, integrate oxytocin and dopamine signaling to render social stimuli rewarding^32,33^. Whether these evolutionarily preserved motivational circuits differentiate face-to-face from online affiliation remains unknown. Resolving this question is critical for understanding whether these circuits primarily support the benefits of close relationships or can be co-opted by social media and emerging technologies in ways that foster excessive engagement and disrupt everyday functioning^8,34^.

Consistent with a shared substrate for dysregulated motivation, overlapping mesocorticolimbic circuits drive substance addiction^35^: drug-associated cues reliably heighten neural reactivity in the ventral striatum and medial frontal cortex across drugs of abuse^36^, with additional dorsal-striatal and insular adaptations promoting habit formation and interoceptive craving. Together with a proposed role for dopamine and oxytocin in addiction^37^ and behavioral similarities encompassing preoccupation, emotional dependence, and craving, these parallels have motivated hypotheses that associate romantic love with addiction^5,6^. Empirical tests, however, have been limited by the absence of a precise neuromarker for romantic love in humans.

Most prior neuroimaging studies of romantic love and other motivational processes have relied on highly controlled paradigms and univariate analyses that excel at localizing peak activation in individual voxels but yield low ecological validity, low-to-moderate effect sizes, and limited sensitivity for comparing the neural representations of complex mental processes^38^. Recent work indicates that combining fMRI with machine-learning-based multivariate pattern analysis (MVPA) provides a more comprehensive and precise approach to modeling highly subjective affective experiences than conventional univariate methods^38–42^. By integrating whole-brain response patterns, MVPA can discriminate distinct brain states with high resolution and sensitivity, and the resulting neurofunctional signatures enable principled tests of the common and separable neural representations of mental processes across domains.

The present project therefore combined naturalistic fMRI with machine-learning-based neural decoding across a series of experiments and datasets (Fig. 1). First, we trained and validated whole-brain neurofunctional decoders of romantic love and friendship and mapped their shared and distinct representations (study 1, *n* = 52). We then tested whether these signatures generalized to social affiliation in the digital domain (study 2, social-media reward, *n* = 54), while remaining distinct from primary and secondary non-social rewards (study 3, gustatory reward, *n* = 57; study 4, monetary reward, *n* = 39). To characterize the affective specificity of the signatures, we further examined their sensitivity to positive versus negative arousing stimuli in two independent datasets (studies 5 and 6, *n* = 60 and *n* = 36). Finally, we applied the resulting models to test whether the love signature is sensitive to the effects of intranasal oxytocin in men viewing their romantic partner versus another woman (study 7, *n* = 40) and whether it captures drug-cue reactivity in heavy cannabis users from non-users (study 8, *n* = 82).

**Fig. 1.**
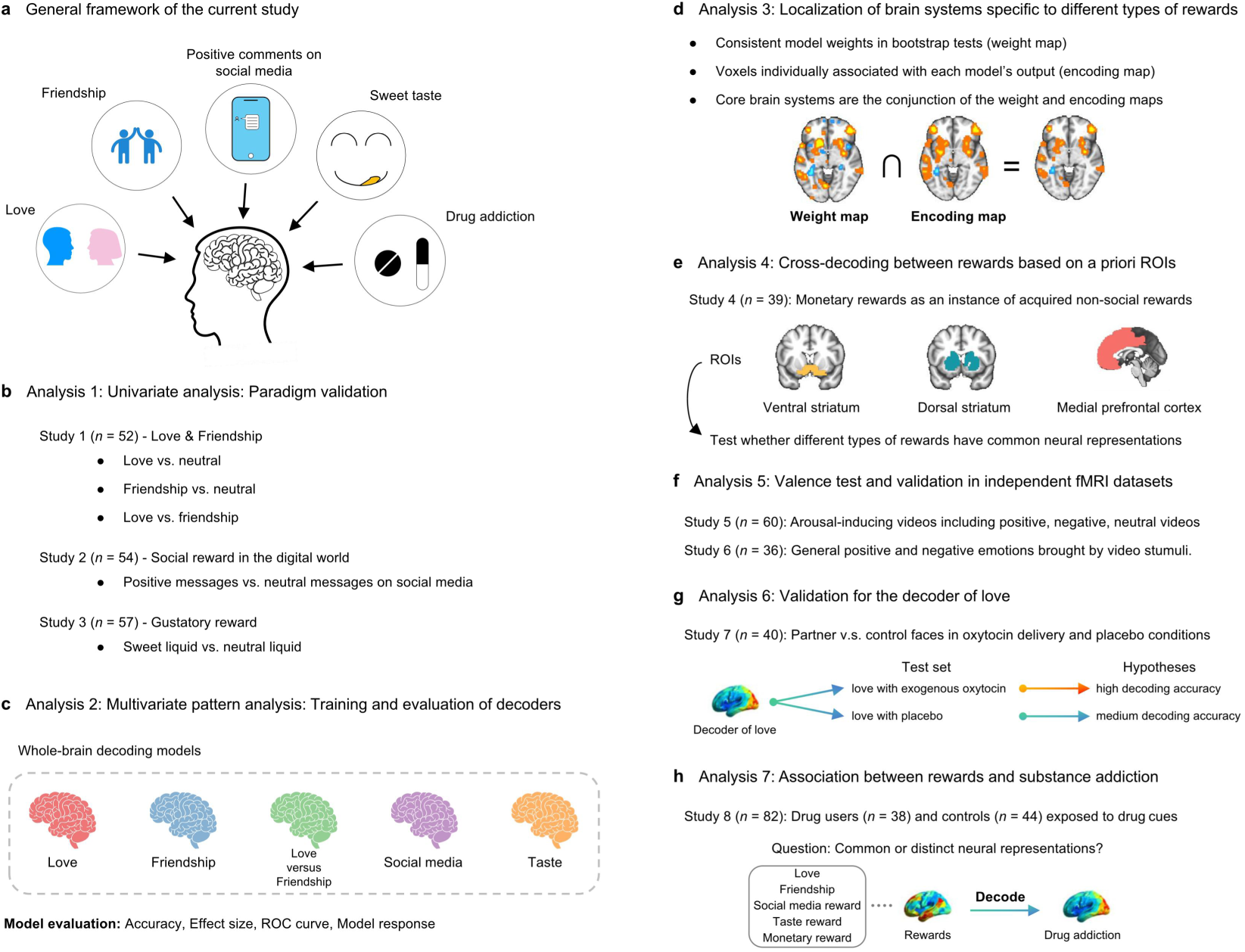
Research framework and main analyses. **a,** The present study combined naturalistic fMRI with predictive modelling to determine shared and distinct neural representations of romantic love and friendship (study 1, *n* = 52), test whether these representations generalize to social affiliation in the digital domain (study 2, social-media reward, *n* = 54) while remaining distinct from non-social rewards (study 3, gustatory reward, *n* = 57; study 4, monetary reward, *n* = 39), and examine whether romantic love and drug addiction exhibit overlapping neural representations (study 8, *n* = 82). **b,** Univariate analysis was employed to preliminarily identify neural circuits involved in social-affiliative and non-social primary rewards. **c,** Decoding model development using support vector machines (SVM) for social-affiliative and non-social rewards. Voxel-level brain maps (beta images) were used as features, and a whole-brain multivariate pattern predictive of rewarding experience was trained. **d,** Identification of core brain regions for each type of reward. Core regions for different types of rewarding experience were defined as the conjunction of the weight map and the model encoding map derived from each decoding model. **e,** Cross-decoding based on regions of interest (ROIs) among different types of rewards was conducted to test whether these rewarding experiences have shared or distinct neural representations. ROIs included ventral striatum, dorsal striatum and medial prefrontal cortex, which were documented as regions engaged in reward processes^43–45^. **f,** Valence tests using positive and negative arousing stimuli (i.e., videos) in additional datasets (studies 5 and 6, *n* = 60 and *n* = 36). **g,** Testing whether the love decoder was sensitive to oxytocinergic modulation of romantic love (study 7, *n* = 40). Particularly, we compared the decoding accuracy levels of the love decoder when decoding brain responses to one’s romantic partner versus an unfamiliar individual in both oxytocin administration and placebo conditions. **h,** Examining the association between romantic love and drug addiction (study 8). The decoder for romantic love was used to decode drug-cue reactivity of heavy cannabis users (*n* = 38) and non-users (*n* = 44). Decoders for other rewards were also included in this classification analysis to test the specificity of the association between romantic love and drug addiction in contrast to other rewarding experiences.

## Results

### Behavioral results

In our primary study (study 1), intense and personally relevant experiences of love and friendship during fMRI were induced by a validated autobiographical memory recall paradigm^46,47^(Fig. 2a). A total of *N* = 52 participants (28 females) who were in an intense romantic relationship and a close cross-sex friendship for longer than six months (Passionate Love Scale scores > 100; McGill Friendship Questionnaire scores > 150; Fig. 3a) underwent recall of their most romantic experiences with their partner, their most pleasant experiences with their friend and neutral experiences from time spent by oneself (yet in a social context, e.g., riding a subway) during MRI acquisition. The neutral experiences served as baseline and involved similar cognitive processes (e.g., episodic memory retrieval, social context) but lacked the specific social affective bonding component present in the love and friendship conditions. Following each memory recall, participants rated their levels of experienced romantic love and pleasantness (ranging from 1 = not at all to 5 = very). Analysis of these ratings confirmed that the paradigm successfully induced strong subjective experiences of romance and pleasantness (Fig. 3a). Reported romantic experience was considerably higher during love memory recall (mean ± SD: 4.55 ± 0.43) compared with both other conditions (friendship vs neutral: *t*(51) = 6.94, Cohen’s *d* = 0.96, *P* < 0.001; love vs neutral: *t*(51) = 51.55, Cohen’s *d* = 7.15, *P* < 0.001; love vs friendship: *t*(51) = 28.85, Cohen’s *d* = 4.00, *P* < 0.001). Both love and friendship conditions induced high levels of pleasantness (mean ± SD; love: 4.68 ± 0.34; friendship: 3.87 ± 0.67), with higher pleasantness in the love condition compared with the friendship condition (friendship vs neutral: *t*(51) = 23.02, Cohen’s *d* = 3.19, *P* < 0.001; love vs neutral: *t*(51) = 44.85, Cohen’s *d* = 6.22, *P* < 0.001; love vs friendship: *t*(51) = 8.73, Cohen’s *d* = 1.21, *P* < 0.001). Therefore, we included pleasantness level as the covariate for later analyses involving comparison between neural representations of love and friendship (e.g., the decoding model of love versus friendship), with findings indicating robustness of the results after controlling for pleasantness level (Supplementary Fig. 2).

**Fig. 2.**
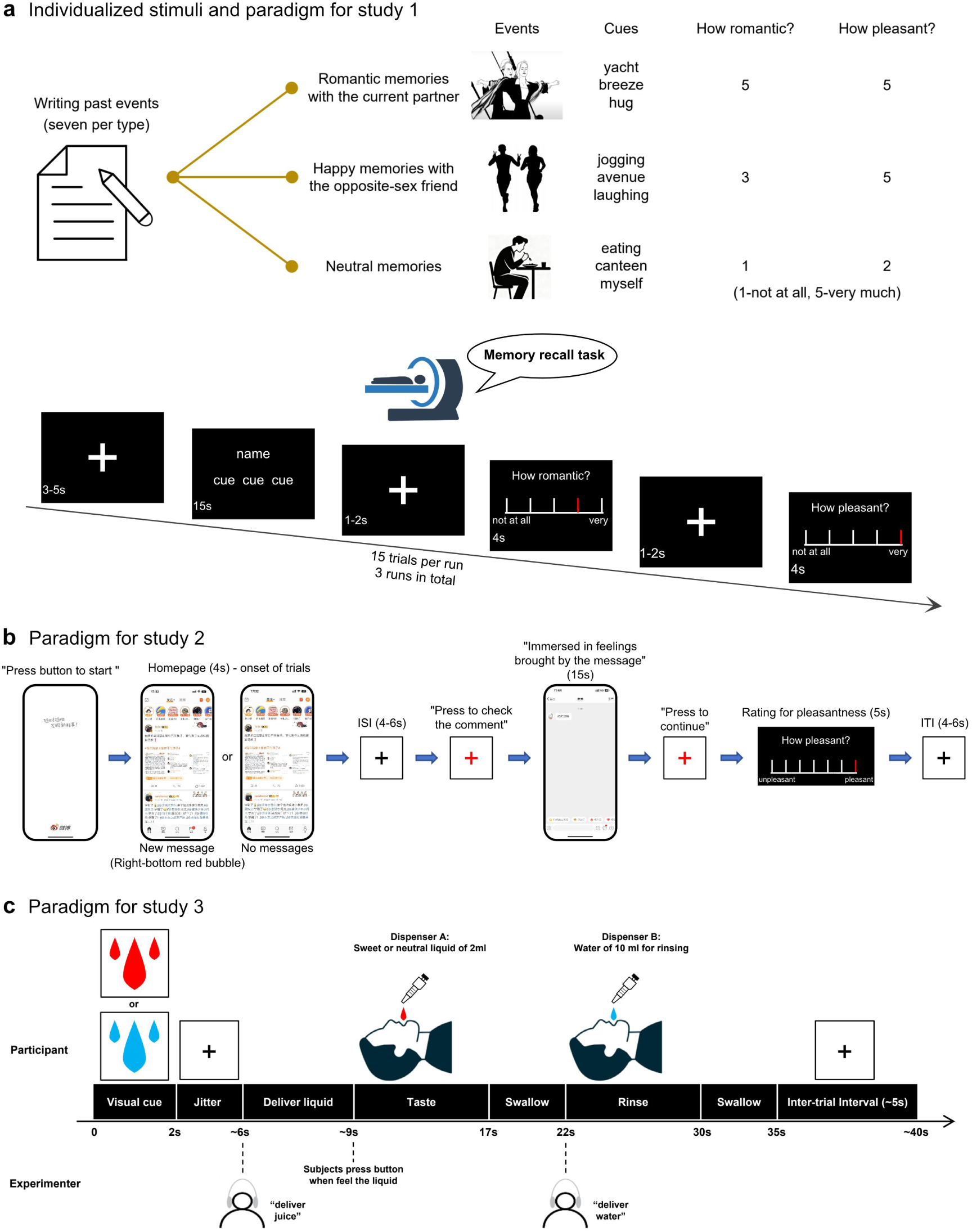
Task designs for studies 1-3. **a,** Procedures for individualizing in-scanner stimuli and the memory recall task used in study 1 (social reward via positive social memories). To determine neural representations of romantic love and friendship, we developed a memory-recall fMRI paradigm in which intense romantic love experience and/or pleasantness were induced by vivid autobiographical memories. In each run of three, participants recalled fifteen past events vividly and re-experienced the romantic love experience brought by their romantic experiences with the current partner, pleasant experiences with a friend from the opposite sex, and neutral experiences from time spent by oneself, before rating levels of experienced romantic love and pleasantness. Neutral experiences served as the baseline and engaged comparable cognitive processes (e.g., episodic memory retrieval, social context) but lacked the social bonding component present in the love and friendship conditions, thereby enabling extraction of relationship-specific representations. **b,** Task design for study 2 (social reward in the digital domain). To examine whether neural representations of real-world social affiliation generalize to social affiliation in the digital world, we designed a naturalistic and personalized experiment during which participants read positive messages that they wanted to receive from close others on social media and neutral messages on simulated interfaces of social media platforms. Then, participants rated levels of pleasantness when reading these messages. **c,** Paradigm for study 3 (taste reward via sweet liquids). To test whether neural representations of social rewards differ from those of non-social rewards, we designed an individualized liquid tasting experiment during which participants received visual instructions and tasted individualized neutral liquids (artificial saliva with different concentrations) or pleasant sweet liquids (milk beverage) five times in each of four runs in the scanner. At the end of each run, participants rated levels of pleasantness when tasting sweet/neutral liquid.

**Fig. 3.**
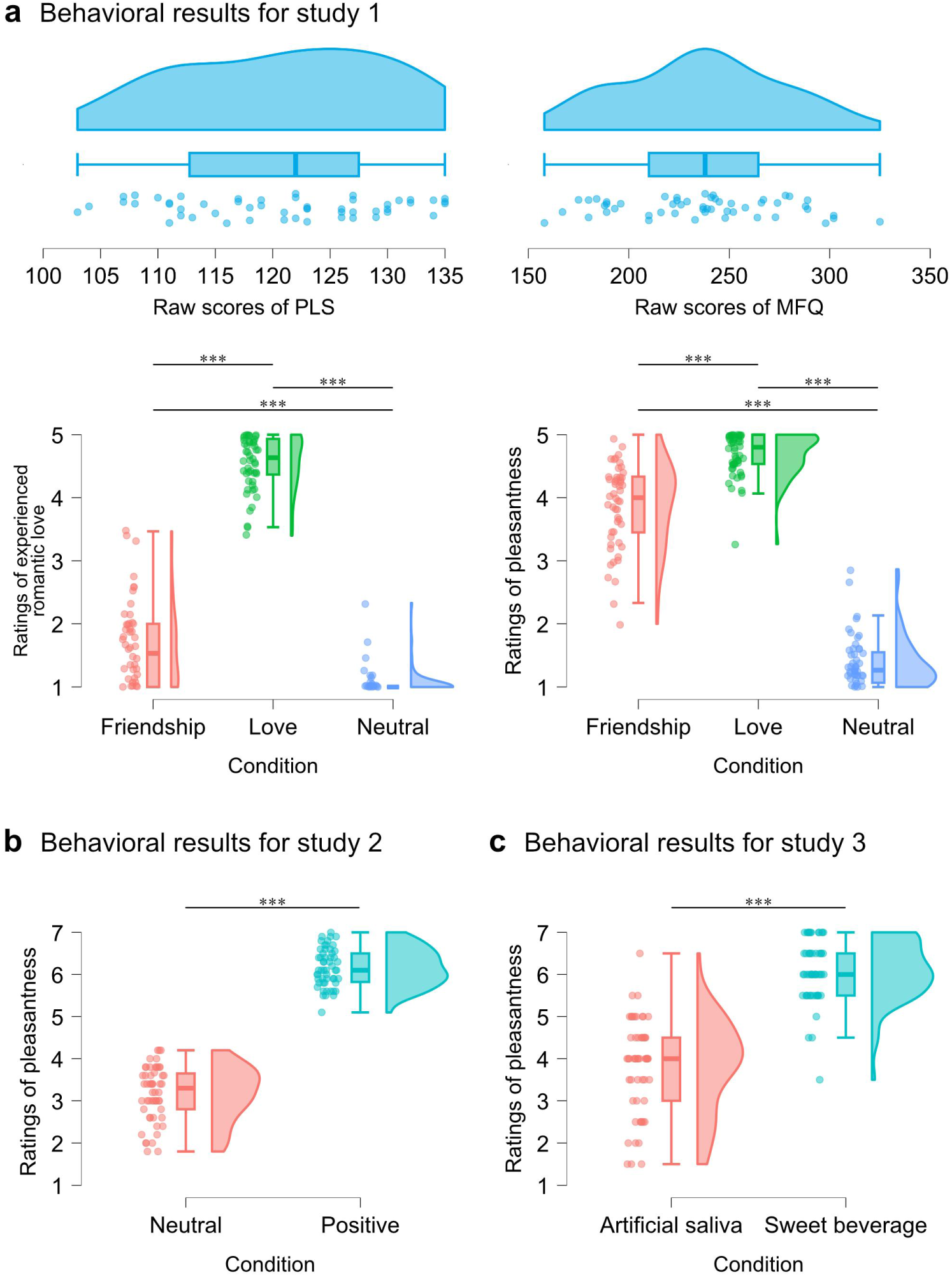
Behavioral results for studies 1-3. The raincloud plots show the distributions of questionnaire scores and behavioral ratings during MRI tasks. **a,** All participants in study 1 reported an intense romantic relationship (Passionate Love Scale score > 100) and a strong cross-sex friendship (McGill Friendship Questionnaire score > 150). Participants reported significantly higher levels of experienced romantic love when recalling romantic memories compared with other conditions and reported higher levels of pleasantness when recalling positive memories with their romantic partner and friend compared with neutral memories. **b,** Participants in study 2 reported significantly higher levels of pleasantness when reading positive comments compared with neutral comments. **c,** The paradigm in study 3 induced significantly higher pleasantness ratings when participants tasted sweet liquids compared with neutral liquids. The boundaries of the box plots represent the first and third quartiles, and the whiskers indicate the range of the median ± 1.5 times the interquartile range. PLS, Passionate Love Scale; MFQ, McGill Friendship Questionnaire. \*\*\**P* < 0.001.

The experiment in study 2 (Fig. 2b) utilized a personalized paradigm that closely resembled positive social experiences on social media platforms and aimed to determine whether positive social experiences in the real and digital contexts can be differentiated. To this end, participants (*n* = 54, 30 females) were shown personalized depictions of their social media accounts while receiving positive comments from close others or neutral comments from bots. Following each event, participants rated their pleasantness (ranging from 1 = not at all to 7 = very). Positive comments from close others elicited strong and significantly higher pleasantness ratings (mean ± SD: 6.16 ± 0.45) compared with neutral comments from bots (*t*(53) = 27.11, Cohen’s *d* = 3.69, *P* < 0.001; Fig. 3b).

The experiment in study 3 (Fig. 2c) implemented a sweet taste paradigm in which participants (*n* = 57, 33 females) were administered sweet or neutral liquids during fMRI (using procedures similar to ref. 40) and subsequently rated the experienced pleasantness (ranging from 1 = not at all to 7 = very). This experiment was designed to distinguish neural signatures of social reward experiences (love and friendship) from those of primary, non-social rewards such as sweet taste. The paradigm robustly induced high levels of subjective pleasantness (mean ± SD for the sweet taste condition: 6.10 ± 0.74), with significantly greater pleasantness reported for sweet compared with neutral liquids (*t*(56) = 15.39, Cohen’s *d* = 2.04, *P* < 0.001; Fig. 3c).

### Univariate evaluation of the paradigms

Univariate analyses were initially conducted to identify neural circuits involved in each type of reward and to provide further validation for the paradigms beyond behavioral ratings. Consistent with prior meta-analytic work^23,48,49^, the striatum and frontal cortex, especially the medial prefrontal cortex, were the most consistently engaged brain regions across different reward types (Extended Data Fig. 1a,b,e,f). In addition, the bilateral precuneus responded to social affiliative rewards from both the real and digital worlds but did not respond to gustatory reward. The bilateral insula was selectively engaged by gustatory reward, whereas the bilateral supplementary motor area was selectively engaged by social reward in the digital domain. Critically, direct comparison of the love and friendship conditions revealed overlapping activation in key nodes of the mesocorticolimbic reward and motivational system, particularly the ventral striatum and medial prefrontal cortex (Extended Data Fig. 1c). The direct contrast further showed that romantic love elicited greater activation in the anterior cingulate cortex, ventral tegmental area, precuneus and bilateral fusiform gyri, whereas friendship elicited greater activation in right lateral prefrontal regions, including the dorsolateral prefrontal cortex and inferior frontal gyrus, as well as the inferior parietal lobule and posterior cingulate cortex (Extended Data Fig. 1d).

### Identifying comprehensive and accurate neural signatures of love and friendship

We employed a support vector machine (SVM, with 10 × 10-fold cross-validation) to develop brain activity-based decoding models for the experiences of love and friendship. The decoders accurately classified both conditions from neutral experiences with high and significant accuracy (love versus neutral: accuracy = 90% ± 4.1% SE, *P* < 10^−6^, Cohen’s *d* = 1.58; friendship versus neutral: accuracy = 94% ± 3.2% SE, *P* < 10^−6^, Cohen’s *d* = 1.73, Fig. 4a,b,c). A direct comparison further indicated that love and friendship could be accurately distinguished (love versus friendship: accuracy = 85% ± 5.0% SE, *P* < 10^−6^, Cohen’s *d* = 1.49, Fig. 4d).

**Fig. 4.**
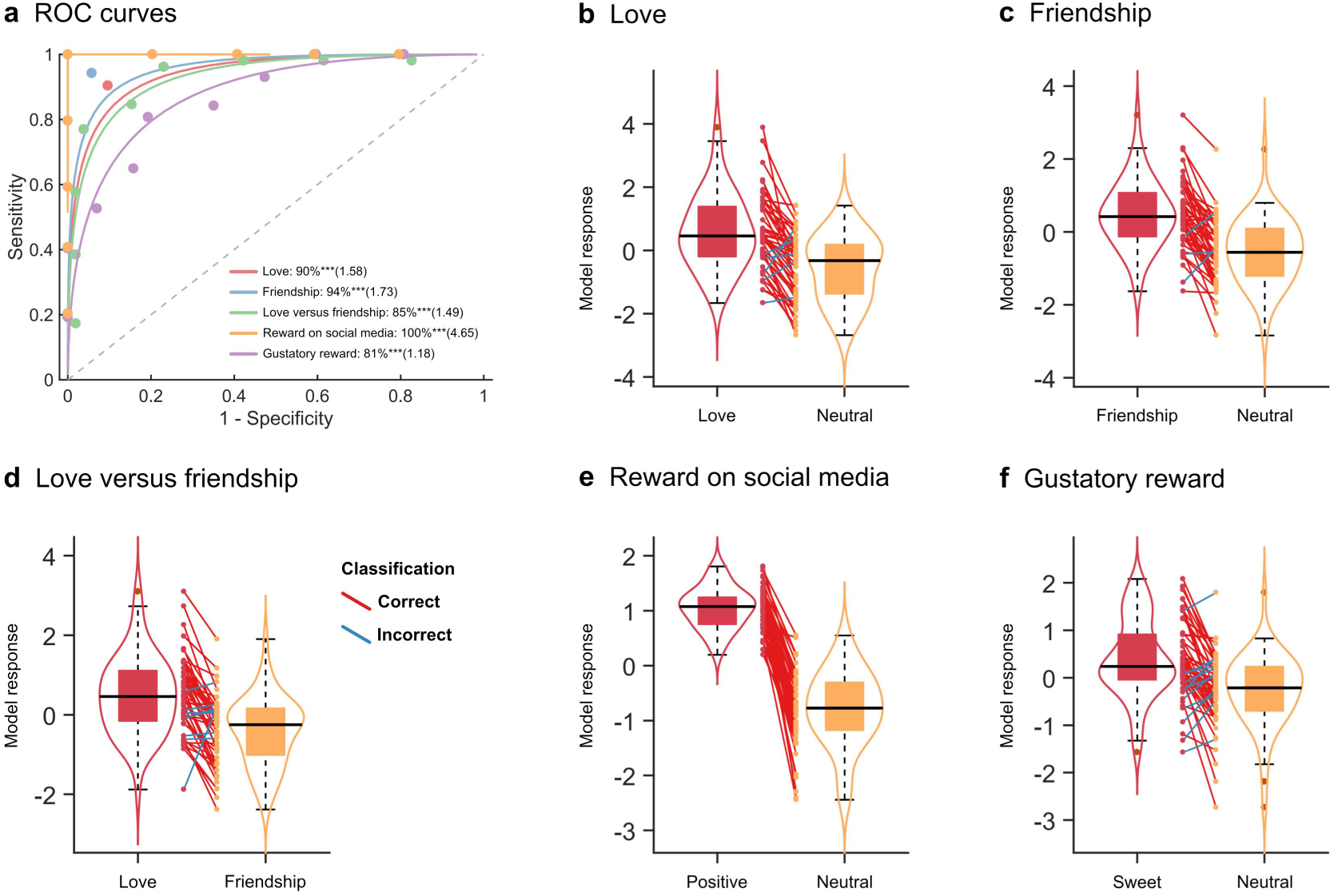
Model performance of whole-brain decoders. **a,** The top-left panel shows receiver operating characteristic (ROC) curves along with forced-choice classification accuracy and Cohen’s *d*. All decoding models for different types of rewarding experiences achieved high accuracy (>80%) as evaluated using 10 × 10-fold cross-validation. **b,c,d,e,f,** Violin plots show model responses for different conditions within each decoder. Each line connecting two dots represents paired data from the same participant (red, correct classification; blue, incorrect classification). The embedded box plots indicate the median (bold horizontal line), the 25th and 75th percentiles (lower and upper hinges), and whiskers mean 1.5 × interquartile range (IQR). \**P* < 0.05, \*\**P* < 0.01, \*\*\**P* < 0.001.

Additional control analysis showed that the decoding accuracy (love versus friendship) remained robust after accounting for pleasantness level (accuracy = 83% ± 5.2% SE, *P* < 10^−5^, Cohen’s *d* = 1.54). The decoding models for reward on social media and sweet taste were also sensitive to their respective rewarding experiences (reward on social media versus neutral: accuracy = 100% ± 0.1% SE, *P* < 10^−10^, Cohen’s *d* = 4.65, Fig. 4e; sweet taste versus neutral: accuracy = 81% ± 5.2% SE, *P* < 10^−5^, Cohen’s *d* = 1.18, Fig. 4f).

To identify robust core brain regions predictive of love and friendship, we combined both model weight maps (Extended Data Fig. 2) and model encoding maps (‘structure coefficients’; Extended Data Fig. 3). Model weights from the corresponding bootstrapping tests identify voxels that reliably contribute to model predictions, while model encoding maps identify voxels whose activity correlates with the predicted response^50^. Their combination therefore facilitates the identification of voxels that are both essential for model prediction and associated with the mental processes under investigation.

Findings from the combined approach revealed that voxels in core regions of the mesocorticolimbic reward and motivational system^43–45^, in particular the nucleus accumbens within the ventral striatum and the medial prefrontal cortex, as well as in the precuneus - a core hub of the default-mode network involved in episodic memory and self-related mental representations^51^-predicted both love and friendship (Fig. 5a,b). Direct comparison of love and friendship indicated that voxels in a more dorsal region of the ventral striatum, namely the putamen, anterior and mid-cingulate regions, and bilateral fusiform gyrus, were more positively predictive of love. In contrast, right lateral prefrontal regions, including the right dorsolateral prefrontal cortex and inferior frontal gyrus, as well as the posterior cingulate cortex, were negatively predictive of love compared with friendship (Fig. 5c).

**Fig. 5.**
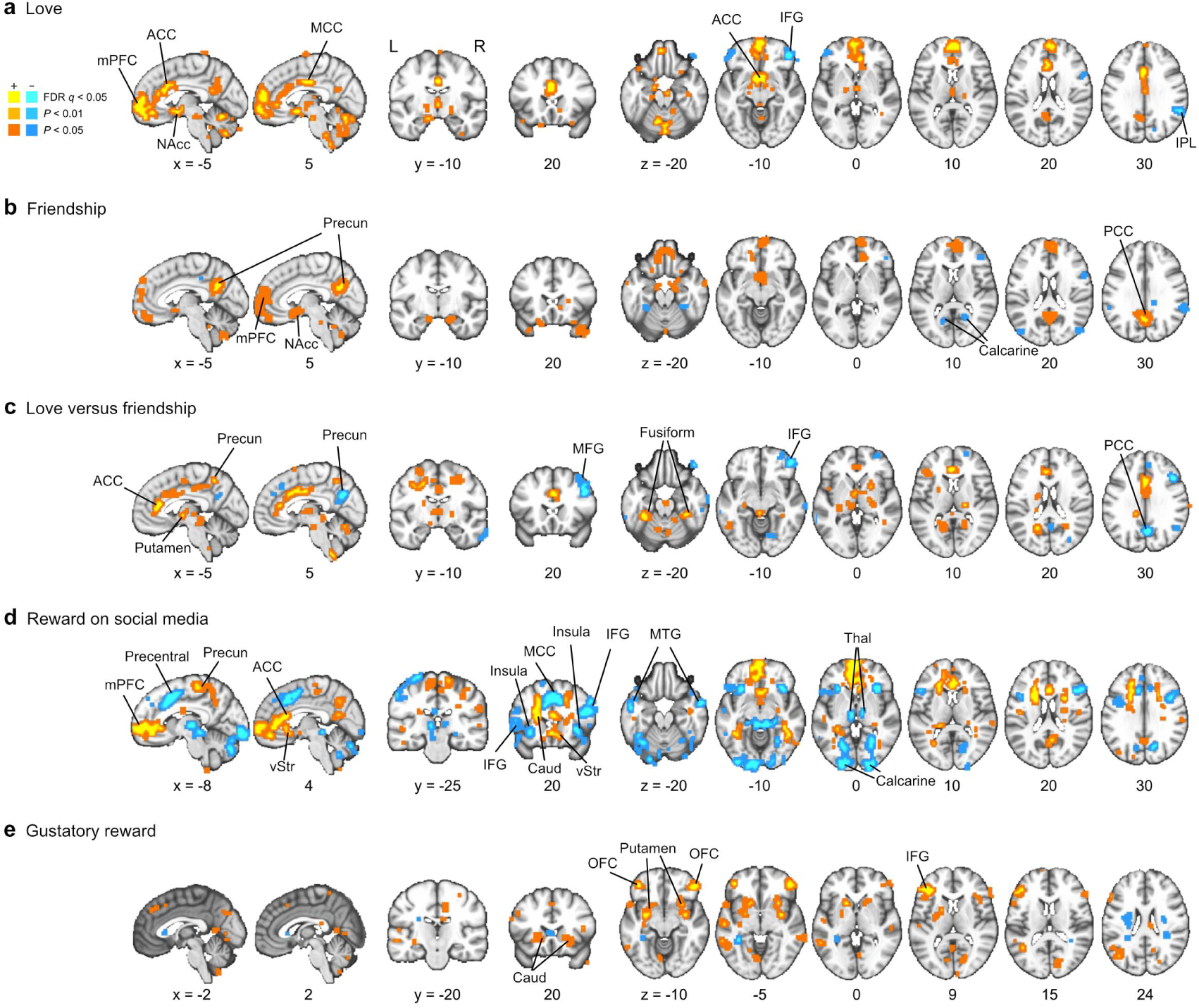
Core brain regions for different types of rewards. **a,b,** Voxels in core nodes of the mesocorticolimbic reward circuitry, particularly the nucleus accumbens within the ventral striatum and the medial prefrontal cortex, together with the precuneus, were jointly predictive of both love and friendship. **c,** A more dorsal sector of the ventral striatum (putamen), anterior and mid-cingulate regions, and bilateral fusiform gyri were more positively predictive of love, whereas right lateral prefrontal cortex (including right dorsolateral prefrontal and inferior frontal regions) and the posterior cingulate cortex carried relatively greater predictive weight for friendship. **d,** Social-media reward was supported by overlapping mesocorticolimbic regions, notably the ventral striatum and medial prefrontal cortex, alongside regions involved in integrating sensory and social information, including the thalamus, supplementary motor area, middle temporal gyrus, and calcarine sulcus. **e,** Gustatory reward was most strongly predicted by regions linked to reward and taste processing, such as the bilateral dorsal striatum and lateral orbitofrontal cortex, as well as primary gustatory areas in the opercular cortex and adjacent inferior frontal and anterior insular cortices. Striatal regions and medial prefrontal cortex were consistently involved in representations of different rewarding experiences. Core regions for different types of rewarding experiences were defined as the conjunction of the weight map and the model encoding map. Multiple thresholds (uncorrected *P* < 0.05 and *P* < 0.01, as well as FDR *q* < 0.05; two-tailed) were employed to illustrate the extent of clusters. mPFC, medial prefrontal cortex; ACC, anterior cingulate cortex; MCC, middle cingulate cortex; Precun, precuneus; vStr, ventral striatum; NAcc, nucleus accumbens; OFC, orbitofrontal cortex; SMA, supplementary motor area; MTG, middle temporal gyrus; Caud, caudate nucleus; IFG, inferior frontal gyrus; Thal, thalamus; IPL, inferior parietal lobe; PCC, posterior cingulate cortex; L, left hemisphere; R, right hemisphere.

The two other reward paradigms were predicted by partly distinct as well as overlapping regions. Reward experience in the social media context was predicted by voxels in the mesocorticolimbic reward system, particularly the ventral striatum and medial prefrontal cortex, as well as a set of regions involved in integrating sensory and social information, including the thalamus, supplementary motor area, middle temporal gyrus, and calcarine sulcus (Fig. 5d). Gustatory reward was preferentially predicted by regions associated with reward processing, such as the bilateral dorsal striatum and lateral orbitofrontal cortex, as well as primary taste areas, including the opercular cortex and adjacent inferior frontal and anterior insular regions (Fig. 5e).

### Comparing neural signatures of romantic love and friendship with other rewarding experiences

Given that social processes, including social reward, may engage both shared and distinct neural pathways^52,53^, and that rewarding experiences may be encoded in a “common neural currency” (e.g., Levy et al.^54^), we next tested whether the neural signatures of social affiliative rewards in the domains of love and friendship are sensitive to a social affiliative reward in the digital world (positive comments on social media, study 2), a non-social primary reward (sweet taste, study 3), and a non-social secondary reward (money, study 4, *n* = 39, see ref. 55). To specifically test the “common neural currency” hypothesis of reward valuation across different modalities and reward targets, we implemented an ROI-based cross-decoding approach^56^, which was restricted to the core reward circuitry including the ventral striatum (VS), dorsal striatum (DS) and extended medial prefrontal cortex (mPFC) (for details see Methods; note that results were consistent with whole-brain decoding models; see also Supplementary Fig. 1). While distributed cortical regions such as the precuneus and posterior cingulate clearly contributed to broader representations of social affiliation (Fig. 5a,b,d), restricting the analyses to a priori striato-frontal reward systems allowed a standardized comparison across social and non-social reward domains while controlling for the different modalities and paradigms. Importantly, the decoders trained on the striato-frontal reward circuits predicted their respective reward types with high accuracy (all *P* < 0.001, effect sizes > 1.08; see diagonal pattern in Fig. 6a,b,c), confirming that these systems are key contributors to decoding across different reward types and paradigms.

**Fig. 6.**
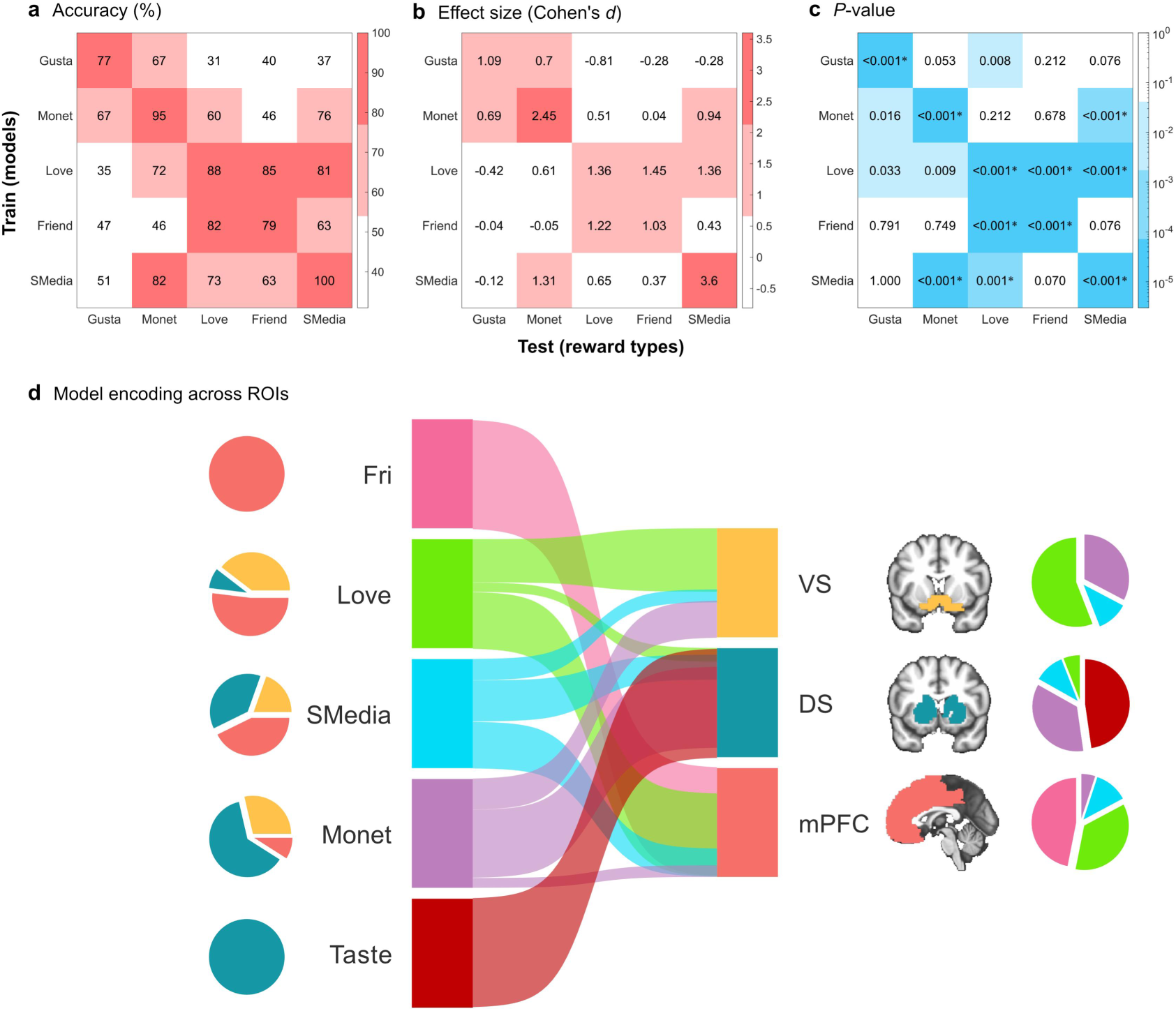
Cross-decoding performance and river plot of decoding models. **a,b,c,** The love decoder significantly cross-predicted friendship and social-media reward, but not gustatory or monetary rewards, suggesting a shared neural basis among social affiliative rewards. In contrast, the friendship decoder did not generalize to social-media or non-social rewards, whereas the social-media reward decoder showed cross-prediction to monetary reward only. Accuracy, effect sizes (Cohen’s *d*), and associated *P*-values were derived from a 10 × 10-fold cross-validation procedure. Values along the diagonal correspond to decoding performance when the model was developed using only voxels within the ROIs. *Bonferroni-corrected *P* < 0.05 (two-tailed). **d,** Interpretation of whole-brain decoding models using river plots. River plots illustrate regional variance across decoders by estimating spatial similarity (cosine similarity) between encoding maps of whole-brain models and a priori ROIs previously identified in neuroimaging studies of reward-related processes^43–45^. Ribbons are normalized to the maximum cosine similarity across all ROIs, and each decoding model is represented by a distinct color. Encoding maps used for computing spatial similarity were thresholded at FDR *q* < 0.05, with only positive voxels retained for similarity calculation and interpretation. Distance between ribbons and boxes is arbitrary and shown for visualization clarity. Pie charts on the left represent the composition of important voxels for each decoding model, whereas those on the right indicate the relative cluster sizes of each ROI across models. Gusta, gustatory; Monet, monetary; Friend, friendship; SMedia, reward on social media; VS, ventral striatum; DS, dorsal striatum; mPFC, medial prefrontal cortex.

Love significantly cross-decoded friendship and reward on social media, but not gustatory or monetary rewards, suggesting a shared neural basis among these social affiliative rewards (Fig. 6a,b,c). Friendship did not cross-predict social media or any non-social rewards, while reward on social media cross-predicted monetary reward. To further examine common and distinct neural representations across reward types and core nodes of the reward circuitry, we generated river plots illustrating the composition of encoding maps derived from whole-brain models. Love primarily engaged the ventral striatum and medial prefrontal cortex, with minimal engagement of the dorsal striatum, whereas friendship predominantly engaged the prefrontal systems (Fig. 6d, left pie charts). Moreover, there was a graded increase in dorsal striatal involvement from love to reward on social media, monetary reward and gustatory reward. Pie charts on the right illustrate the relative sizes of suprathreshold clusters within each ROI across whole-brain models.

### Contributions of nonspecific arousal to decoding performance

Given that nonspecific affective arousal contributes to the performance of neurofunctional decoders^38^, we next tested the model response on fMRI data acquired during highly arousing positive and negative movie clips. To rule out this explanation, we examined whether our models showed preferential responses to positive versus negative stimuli in two additional datasets (study 5, *n* = 60; study 6, *n* = 36). After examining model responses in positive minus neutral and negative minus neutral stimuli respectively, we found that all whole-brain decoding models responded significantly to highly arousing positive stimuli (Cohen’s *d* > 0.86) but showed non-significant responses to highly arousing negative stimuli (Cohen’s *d* < 0.18), indicating specificity for positively valenced stimuli (Extended Data Fig. 4).

### Sensitivity of the decoders to oxytocin-enhanced love experience

Oxytocin plays a pivotal role in monogamous partner bonding across species^2^, and previous pharmacological imaging studies in humans have shown that intranasal oxytocin enhances reward-system responses and perceived attractiveness in men when viewing the face of their romantic partner, but not that of an unfamiliar woman^14^. We leveraged data from this human study (study 7, *n* = 40 males), in which participants received oxytocin (24 IU) and placebo nasal spray in a within-subject, placebo-controlled pharmacological fMRI design, to determine the sensitivity of our love decoder to detect an oxytocin-induced enhancement of love (for similar rationale, see ref. 57) and also to provide a more precise test of oxytocin’s role in this domain. Based on neural activity, the love decoder accurately predicted whether men viewed the face of their romantic partner versus a control condition (unknown woman) following oxytocin (accuracy = 83% ± 6.0% SE, *P* < 0.001, Cohen’s *d* = 1.34), but not following placebo (accuracy = 57% ± 7.8% SE, *P* = 0.430, Cohen’s *d* = 0.30), with a significant difference between conditions (*Z* = 2.440, *P* = 0.015; Fig. 7a). In contrast, the friendship decoder could not predict whether the face of the romantic partner or another face was shown under either oxytocin (accuracy = 65% ± 7.5% SE, *P* = 0.081, Cohen’s *d* = 0.19) or placebo (accuracy = 55% ± 7.9% SE, *P* = 0.636, Cohen’s *d* = 0.40), and the difference in accuracy between conditions was not significant (*Z* = 0.913, *P* = 0.361; Fig. 7b). Together, these findings validate the specificity of the decoders for their respective target emotions (love and friendship), while also underscoring the sensitivity of the love decoder to experimentally induced enhancement of romantic love and highlighting the role of oxytocin in monogamous bonding.

**Fig. 7.**
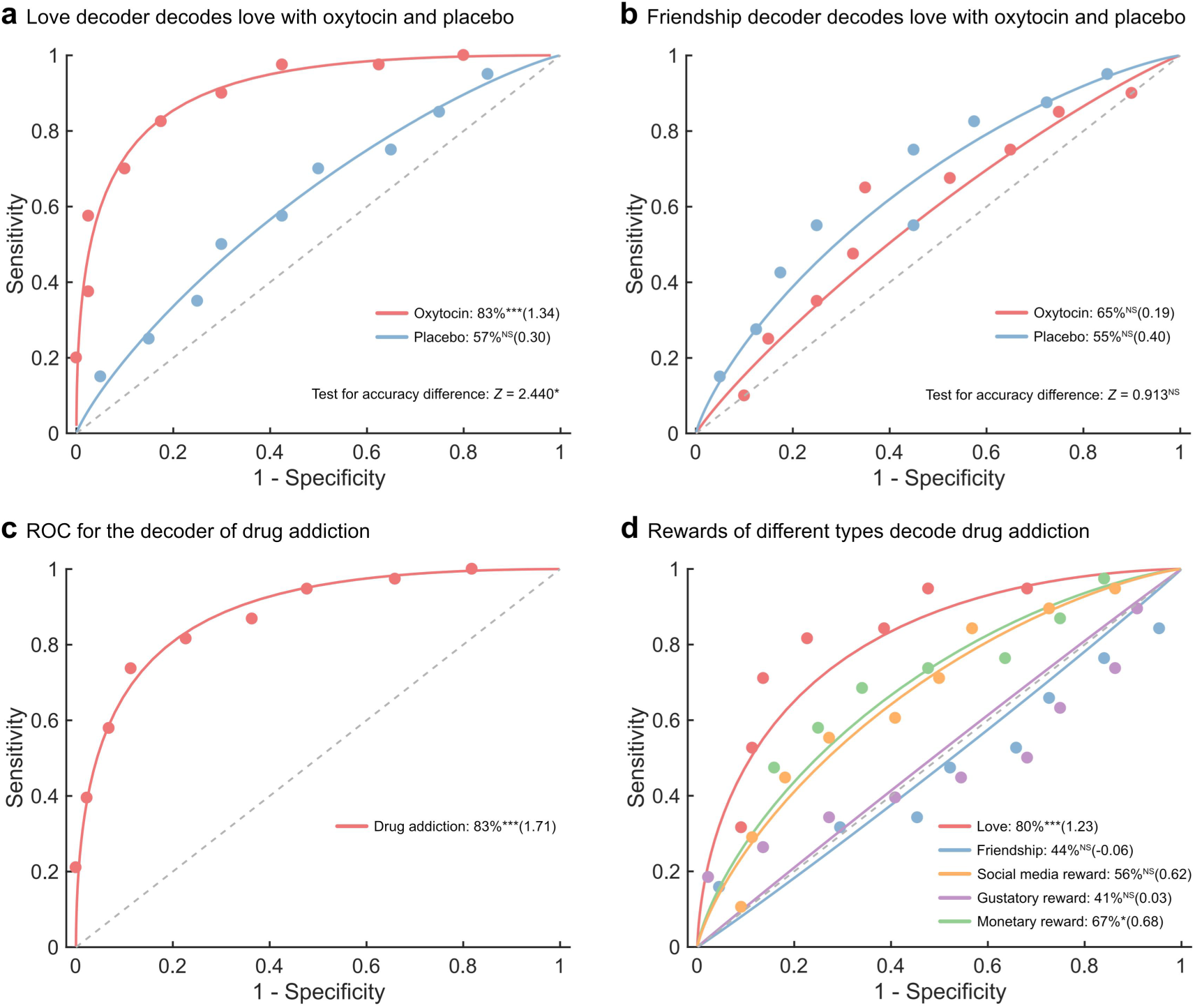
Results for classification analyses. **a,** Receiver operating characteristic (ROC) curves for love experiences in oxytocin and placebo conditions decoded by the love decoder using forced-choice classification (restricted to reward-system ROIs). Decoding accuracy following oxytocin administration was significantly higher than that in the placebo condition. **b,** ROC curves for love experiences in oxytocin and placebo conditions decoded by the friendship decoder using forced-choice classification. The friendship decoder could not capture the effect of oxytocin on love experiences. **c,** ROC curve for the decoding model (restricted to reward-system ROIs) developed using the drug addiction dataset, with single-interval classification accuracy and Cohen’s *d*. **d,** Receiver operating characteristic (ROC) curves for reward decoding in drug addiction (restricted to reward-system ROIs). The love decoder accurately distinguished brain responses to visual drug cues in heavy cannabis users versus non-users. \**P* < 0.05, \*\**P* < 0.01, \*\*\**P* < 0.001, NS not significant.

### Translation into clinical application: love and drug addiction

Previous frameworks and studies have proposed a neurofunctional similarity between love and drug addiction; however, direct neurobiological evidence remains limited, and these hypotheses remain debated^5,6,58^. To explore the neurofunctional similarity between love and addiction, and to enhance the translational potential of the decoding models^42,59^, we tested the decoder for love, restricted to reward-system ROIs (ventral striatum, dorsal striatum, and extended medial prefrontal cortex), on the neural responses to visual drug cues in heavy cannabis users and non-users. This paradigm, referred to as drug-cue reactivity, induces robust activation in reward-related regions in drug users but not in non-users^60^. Testing the decoders on cue reactivity data from a sample of *n* = 38 of cannabis users and *n* = 44 of non-users^36^ (study 8) revealed that the love decoder accurately distinguished cannabis users from non-users (accuracy = 80% ± 4.4% SE, *P* < 10^−4^, Cohen’s *d* = 1.23; Fig. 7d), with accuracy comparable to that of a drug addiction decoder trained directly on the addiction data (accuracy = 83%± 4.2% SE, *P* < 10^−5^, Cohen’s *d* = 1.71; Fig. 7c and Supplementary Fig. 3). In contrast, the friendship decoder, as well as decoders for gustatory reward and social media reward, did not accurately capture drug-cue reactivity (accuracy range: 41% - 56%). A decoder trained on monetary reward could distinguish cannabis users from non-users, but with lower accuracy and effect size (accuracy = 67% ± 5.2% SE, *P* = 0.019, Cohen’s *d* = 0.68; Fig. 7d). Moreover, spatial similarity analyses between unthresholded weight maps of the love decoder of the present study and a drug craving decoder from a recent study^59^, as well as the decoder of drug-cue reactivity trained on data from study 8, revealed similar patterns of positive and negative weights (Fig. 8a,b). Together, these findings provide evidence for a neurofunctional similarity between romantic love - but not friendship - and drug addiction, and underscore the clinical translational potential of neurofunctional decoders.

**Fig. 8.**
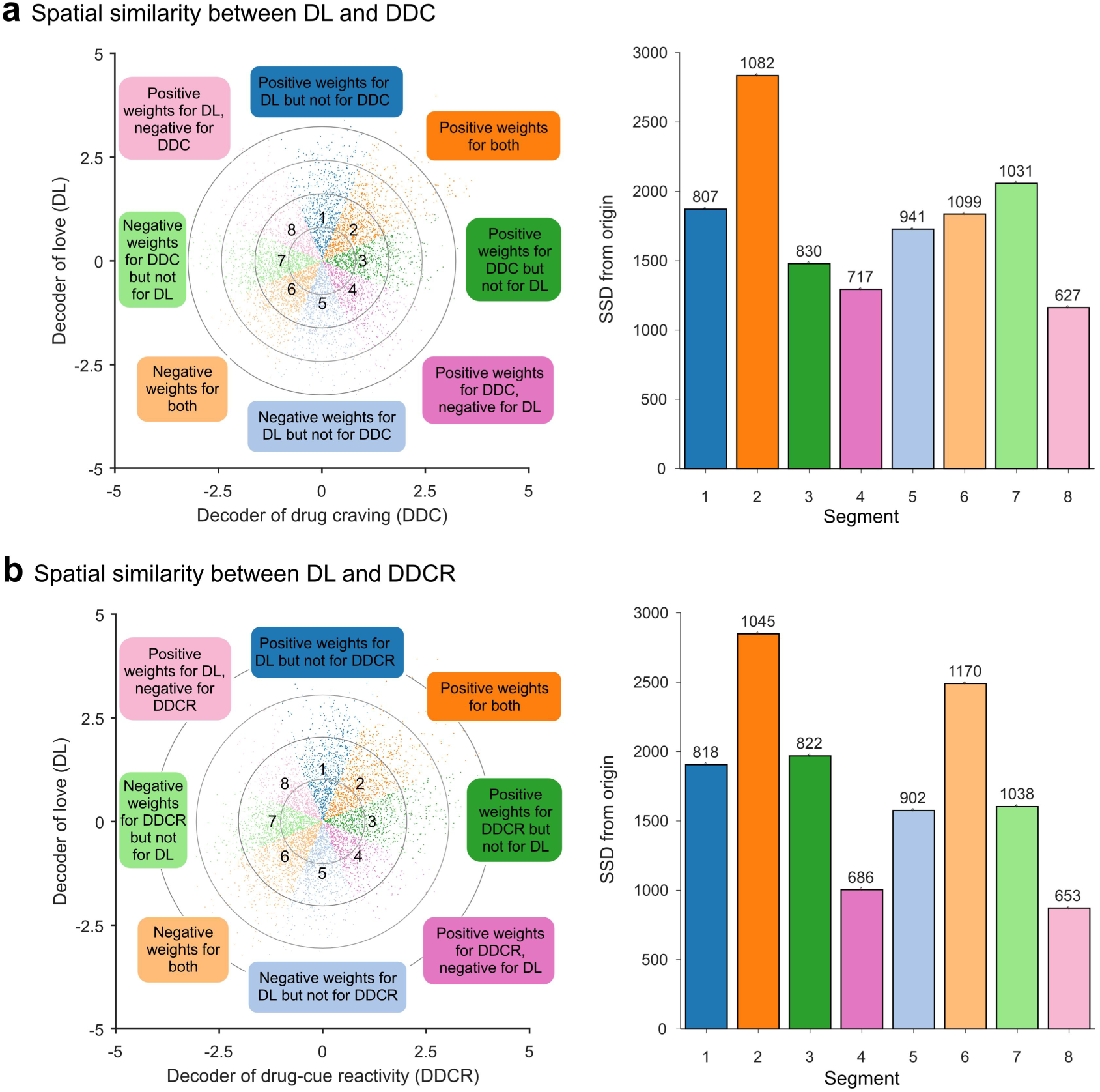
Voxel-level spatial similarity between decoders. The weight maps for the decoder of love (DL) in the present study, the decoder of drug craving (DDC)^59^, and the decoder of drug-cue reactivity (DDCR)^36^ were compared within a mask including the mPFC, ventral striatum, and dorsal striatum. **a,** Spatial similarity analysis between DL and DDC. **b,** Spatial similarity analysis between DL and DDCR. DL and DDC, as well as DL and DDCR, showed similar patterns of positive and negative weights, as the number of voxels exhibiting consistent weights (i.e., positive in both decoders or negative in both decoders) was greater than other combinations. The scatter plot on the left displays normalized weights for two decoders. The radii of the four gray circles represent scaling factors, which are computed as 0.2×, 0.4×, 0.6×, and 0.8× of the maximum weight value in the scatter plot. Each octant corresponds to a specific color, indicating the sign of weights for the two decoders (e.g., shared positive weights are shown in Segment 2, and shared negative weights in Segment 6). The bar plots on the right represent the sum of squared distances from the origin (0,0) for each octant, indicating the overall magnitude of weights in each octant. The number of voxels in each octant is shown above each bar.

## Discussion

Romantic love is among the most powerful human motivational drives, distinguished from broader affiliative functions by its intense and highly selective focus on a single individual. Animal models have established that the formation of selective monogamous bonds depends critically on the dopaminergic mesocorticolimbic reward circuitry - which also mediates broader reward and affiliative processes - and on its interaction with oxytocin^3^. Yet, in contrast to this detailed mechanistic understanding in animal models, a comprehensive brain-wide neurofunctional architecture of the subjective experience of romantic love in humans - its differentiation from friendship, its modulation by oxytocin, and its relationship to addiction - has remained unestablished^1,61,62^.

Here, we combined a series of naturalistic fMRI experiments with machine-learning-based predictive modelling and pharmacological and addiction datasets to determine and comprehensively evaluate the neural representation of the romantic-love experience induced by vivid autobiographical memories. We found that love and friendship jointly engaged the nucleus accumbens, medial prefrontal cortex and precuneus (Extended Data Fig. 1c), yet the two affiliative states were reliably distinguished at the whole-brain level (Fig. 4d) and showed differential engagement of social and cognitive brain systems (Fig. 5c). Given previous theories suggesting that rewarding experiences, including affiliation, may be encoded as a “common neural currency”^52–54^, we further examined the generalizability of the love signature. It accurately cross-decoded the neural expressions of friendship and reward on social media, but not gustatory or monetary reward, indicating a shared neural basis for social-affiliative rewards across the real and digital worlds (Fig. 6a,b,c). Control analyses confirmed that the love decoder was sensitive to positively but not negatively valenced experience, arguing against a strong confounding contribution of affective arousal to decoding performance (Extended Data Fig. 4). We then applied the affiliation-sensitive decoders to show that a single intranasal dose of oxytocin significantly enhanced the neurofunctional discriminability of the romantic partner from another person via the love signature in the reward circuits (Fig. 7a), whereas oxytocin did not affect sensitivity to the neural representation of friendship (Fig. 7b). Finally, we demonstrated that the love decoder - but not the decoders for friendship, social media reward and gustatory reward - accurately detected neural responses to drug cues and distinguished drug users from non-users on the basis of this neurofunctional reactivity (Fig. 7d). Together, these findings reveal a common mesocorticolimbic architecture underlying affiliative states across the real and online worlds, yet distinct from non-social rewards. Within this shared architecture, romantic love is characterized by its additional recruitment of social-cognitive systems and by its selective sensitivity to oxytocin and drug-cue reactivity - offering neurofunctional support for theories of love and its relation to oxytocin and addiction.

The autobiographical, naturalistic paradigms induced vivid and intense subjective experiences of romantic love and pleasure. Consistent with recent meta-analytic evidence, romantic love and friendship engaged partially overlapping representations spanning core reward and motivational regions (nucleus accumbens and medial prefrontal cortex) and regions supporting self-referential processing and autobiographical memory (precuneus)^23^ - together encoding the shared experiences and mutual familiarity that accumulate over time^22,63^. In line with recent work showing that subjective affective states can be accurately decoded from distributed brain activity^38,45,64^, both love and friendship were reliably decodable from neutral experiences based on distributed brain-wide representations. Yet the two neural expressions were distinguishable at the whole-brain level and, at the regional system level, engaged distinct systems that may track attachment-specific information: love more strongly recruited dopaminergic regions implicated in reward and habit formation (ventral tegmental area, putamen) and regions supporting empathy and person-specific identity (anterior cingulate and fusiform gyrus)^65–67^, while compared to friendship, showing weaker engagement of lateral prefrontal regions involved in cognitive control^68^ and of posterior cingulate cortex involved in mentalizing^69^. These findings remained robust after accounting for differences in experienced pleasantness (Supplementary Fig. 2c and Fig. 5c). This dissociation indicates that love links a core reward and motivational circuitry to systems supporting selective attachment to a specific person, whereas friendship additionally engages systems involved in planning and self-other integration.

Bidirectional cross-decoding patterns showed that the love signature tracked friendship and social rewards in the online world - but not sweet taste or monetary reward - further underscoring a core social-reward and affiliation system that is partly distinguishable from primary and secondary non-social rewards. Notably, the neural signature of love extended to a paradigm simulating personalized social-media experiences, in which participants received positive feedback. This converges with prior neuroimaging evidence that receiving online social approval (e.g., “likes” or positive comments) engages core reward circuitry - including the ventral tegmental area, striatal regions, and medial frontal and orbitofrontal cortices - and elicits subjective reward^28–31,70,71^. Meanwhile, we found that the friendship signature did not accurately capture reward on social media. This asymmetry may indicate that digital social rewards preferentially engage highly salient motivational reward systems that overlap with neural representations of romantic love. By contrast, friendship-related representations were more associated with precuneus and posterior cingulate cortex supporting self-other integration and perspective-taking, as well as lateral prefrontal regions reflecting a greater contribution of cognitive control and long-term social cooperation characteristic of real-world friendships (Fig. 5a,b,d and Fig. 6d).

Furthermore, the similarity between representations of romantic love and online social reward may suggest that digitally mediated social rewards may recruit the same neurofunctional circuits that drive real-world affiliation and attachment, and may therefore partly substitute for offline attachment and social support, while also providing a substrate for excessive engagement. This converges with initial accounts proposing that evolutionarily shaped social brain circuits may engage with emerging technologies^7,8^, with inherent opportunities and challenges. On the one hand, digital platforms may be harnessed to establish and sustain social affiliation^72–75^, particularly when direct social contact is not possible or for some individuals who may favor them for their immediate feedback and accessibility^76,77^ or because they lower the barrier imposed by social anxiety^78,79^. On the other hand, a subset of vulnerable individuals may develop problematic, excessive use that, through neuroplastic adaptations in motivational and reward pathways, can escalate toward compulsive engagement and potentially detrimental consequences^80,81^.

Taken together, the finding that real-world and online social rewards are encoded in similar brain representations suggests that the evolutionarily ancient systems supporting social affiliation do not categorically distinguish real-world from digital engagement. This offers an opportunity for social media to act as a scaffolding pathway to affiliation - delivering a neural approximation of real-world attachment and social support in contexts where in-person interaction is constrained, while at the same time presenting a vulnerability to potentially harmful patterns of engagement.

Animal models have underscored the role of oxytocin in highly selective pair bonds^2,9–11^, and a previous human study showed that a single intranasal dose enhances nucleus accumbens responses specifically when men view their partner rather than an unfamiliar woman^14^. Capitalizing on this dataset, we found that the love decoder - but not the friendship decoder - was sensitive to oxytocin, which enhanced the neurofunctional differentiation of the romantic partner but not for others. These findings emphasize that oxytocin does not act on social information in general^82,83^ but on a specific, distributed representation of the bonded partner, and bridge the role of oxytocin from animal models of pair-bonding to the human subjective experience of romantic love - indicating an evolutionarily conserved mechanism of partner bonding^2,9–11^ that extends to the subjective experience of complex human emotion such as love.

Beyond its adaptive role in forming lasting bonds with positive effects on mental health, the engagement of these circuits may also underlie compulsive and ultimately harmful behaviors that extend beyond social media use. Several accounts have proposed a behavioral and neural link between romantic love and drug addiction, motivated by their shared behavioral features - preoccupation, heightened salience of the target, and withdrawal-like states following separation or relationship breakdown^5,6^. Yet, despite initial supporting evidence from a recent meta-analysis^58^, a direct association between love and addiction has remained difficult to establish. Here, we found that the romantic-love signature - but not the signatures for friendship, social media reward and gustatory reward - accurately decoded brain-activity patterns of drug-cue reactivity in heavy cannabis users from non-users (Fig. 7d), indicating that romantic love and drug addiction share neural representations within the reward system. In line with theoretical accounts, this may reflect that addictive substances co-opt neural circuits that originally evolved to mediate selective attachment to a romantic partner^5,6^. Converging support comes from a recent study in which smokers viewing images of their romantic partner together with smoking cues showed reduced cigarette cue reactivity and increased striatal activation relative to viewing an acquaintance with smoking cues^84^. These parallels carry translational implications: if romantic attachment and drug addiction recruit overlapping reward representations, the motivational salience of a partner bond could be harnessed therapeutically - for example, in attachment-based or partner cue-based interventions that compete with or attenuate drug-cue reactivity, as the smoking-cue findings suggest^84^.

More broadly, a shared-circuit account positions the social-bonding system, and its oxytocinergic modulation, as a candidate target for addiction treatment, reframing healthy social attachment not merely as a correlate of recovery but as a potential mechanism of it. The present findings may also provide mechanistic support for an increasing number of studies suggesting a therapeutic potential of oxytocin for addictive disorders^85^, with recent meta-analytic data indicating that oxytocin can alleviate withdrawal symptoms, negative affective states, craving for addictive substances, and substance use in patients^86,87^ and that intranasal administration of oxytocin could help reduce drug-seeking behavior and substance use^88^.

While our study provides a comprehensive neurofunctional model of romantic love, several questions remain open. First, our focus on short-term romantic relationships leaves the neural representation of long-term bonds, such as established companion love, unexplored. The intensity of romantic love typically peaks within the first twelve months, whereas behavioral indicators of a robust attachment bond, such as the use of the partner as a secure base, often emerge only after two or more years^89^.

Future studies should therefore examine how relationship duration is encoded in the brain and how it modulates the overall neural representation of social bonds. Second, our design relied on autobiographical-memory-induced romantic experience.

Although this reliably evoked a strong and vivid experience of romantic love, it does not capture the dynamic temporal structure of the recalled episodes; it is therefore possible that periods of pleasure and of craving for connection were interleaved within a given recall period. Finally, the present study did not explicitly examine the role of attachment style or sex, which may shape the neural representation of love and its modulation by oxytocin^90,91^.

In conclusion, the present study established a comprehensive neurofunctional architecture of the subjective experience of romantic love. This architecture distinguishes romantic love from friendship and from primary and secondary non-social rewards, while revealing shared representations across real-world and digital affiliative rewards. The love signature was specific to positive valence, sensitive to oxytocin during exposure to pictures of a romantic partner, and captured drug-cue reactivity in heavy cannabis users. These findings provide a brain-wide neurofunctional marker of romantic love and offer direct support for theoretical accounts proposing that human pair-bonding relies on evolutionarily conserved, oxytocin-sensitive reward systems that can also be engaged by addictive substances.

## Methods

### Overview

Data from a total of eight studies were used in this paper (Fig. 1 and Supplementary Table 1). Study 1 (*n* = 52), study 2 (*n* = 54) and study 3 (*n* = 57) were originally designed and implemented by the authors, and were used for decoding model development and all main analyses. Studies 4-8 involved secondary analyses of anonymized data from previously published papers^14,36,38,55^. Study 4 (*n* = 39) employed monetary reward stimuli serving as an example of acquired (secondary) non-social rewards to test whether neural signatures of social reward generalize to non-social secondary rewards. Study 5 (*n* = 60) and study 6 (*n* = 36) employed highly arousing positive and negative as well as neutral control video clips and served to test for valence and arousal effects. Study 7 (*n* = 40) employed a romantic love paradigm during which healthy males were exposed to pictures of their romantic female partner or an opposite-sex stranger following the intranasal administration of placebo or oxytocin, and we tested whether the love decoder could detect oxytocin-induced enhancement of neural responses associated with romantic love. Study 8 (*n* = 82) tested neural drug-cue reactivity by exposing cannabis users and non-using controls to visual drug cues and was used to test whether the social reward signatures generalize to addiction-related ‘wanting’. Study 1 was approved by the ethics committee of the University of Electronic Science and Technology of China (Approval number: 1061423061626067). Studies 2 and 3 were approved by the Human Research Ethics Committee of the University of Hong Kong (Approval number: EA240433). Informed consent was obtained from all participants before each of these experiments. For studies 4-8, detailed descriptions of the ethics approval are available via the corresponding references.

### Participants

Participants for the original studies (studies 1-3) were healthy individuals aged between 18 to 30, with the following exclusion criteria: 1) current or history of a physical or mental disorder; 2) drug addiction or substance addiction, or regular and current use of psychoactive substances including cigarettes, e-cigarettes and medication; 3) severe allergies, in particular food allergies; 4) having tattoos, metal dentures, or metal implants in the body; 5) having claustrophobia; 6) other exclusion criteria for MRI acquisition.

Additional inclusion criteria for study 1 (social rewards from positive social memories) were: 1) currently engaged in a committed and exclusive romantic relationship; 2) having a close friendship with a person from the opposite sex; 3) both romantic relationship and cross-sex friendship have been lasting longer than six months; 4) intense romantic relationship (score of passionate love scale > 100); 5) high-quality friendship with the cross-sex friend (score of McGill friendship questionnaire > 150); 6) meeting the romantic partner in person at least once a week; 7) not married; 8) childless. Two participants were excluded from subsequent analyses because of excessive head motion during fMRI scanning (> 3 mm translation or > 3 degrees rotation), leading to a final sample size of *n* = 52 for study 1 (28 females; mean ± SD of age in year: 20.16 ± 1.83; score of passionate love scale: 121 ± 9.0, range: [103, 135]; score of McGill friendship questionnaire: 236 ± 39, range: [158, 325], also see Fig. 3a for data distribution).

For study 2 (social rewards in the digital domain), participants were required to meet the following inclusion criteria: 1) active users of TikTok, Weibo, Red Note or Instagram; 2) engagement with social media for more than four days a week; 3) duration of each social media engagement exceeds 15 minutes; 4) having the same frequently-used social media app as their friends/family do; 5) a rating higher than 4 for the question “To what extent do you believe you can receive messages from friends/family?” (1-do not believe at all, 7-firmly believe). Data of three participants were excluded due to excessive head movement, leading to a final sample size of *n* = 54 (30 females; mean ± SD of age: 20.73 ± 2.17 years).

No additional recruitment criteria were applied for study 3 (gustatory rewards via sweet liquids). Data of five participants were discarded due to excessive head movement, and the final sample size was *n* = 57 (33 females; mean ± SD of age: 19.82 ± 1.59 years).

### Stimuli and paradigm

In study 1 (social reward via positive social memories; Fig. 2a), participants first wrote down seven past events in detail (including location, date, context, process, feelings) for three categories respectively: romantic experiences with the current partner (love condition), pleasant experiences with a friend from the opposite sex (friendship condition) and neutral experiences from time spent by oneself (neutral condition). For each event, participants were required to provide three keywords as cues facilitating a quick and vivid memory recall (e.g., name of their partner, occasion, place). Participants next rated levels of experienced romantic love and pleasantness for each event on 5-point Likert scales (1-not at all, 5-very). Five out of seven events per condition were used in the subsequent fMRI experiment, and the remaining events were used to test whether participants were able to recall the events rapidly. The fMRI experiment was scheduled on a separate day, and only participants who provided vivid and clear descriptions of the events in response to test cues underwent the later fMRI experiment. Participants were required to complete the autobiographical memory recall of the event during the fMRI acquisition (Fig. 2a). Each trial started with a fixation period of 3-5 seconds, followed by the three cue words indicating that participants were required to recall the cued event vividly and re-experience the feelings experienced during this event during the next 15 seconds. After that, participants rated levels of experienced romantic love and pleasantness successively on a rating scale ranging from 1-Not at all to 5-Very strong. Both ratings were preceded by a central fixation cross with a jittered duration of 1-2 seconds. The order of the two ratings was counterbalanced between subjects, and the order of the presented cues of 15 events was randomized. Participants underwent three runs in total, with 15 trials for each. The five trials per condition used in run 1 were repeated in run 2 and run 3 with newly randomized orders to balance memory load versus habituation.

Study 2 (social reward in the digital world; Fig. 2b) was designed to determine the neural basis of social rewards on social media and served to test whether the social reward signatures generalize to rewarding experiences in digital contexts. We designed a naturalistic and personalized experiment during which participants received social rewards (e.g., positive messages, compliments, encouragement, etc.) or non-reward (neutral control messages from a bot). To facilitate a naturalistic and immersive engagement, participants were requested to provide the experimenters with screenshots of their account on their favorite social media platform (e.g., Instagram) prior to the MRI experiment to design individualized stimuli. In addition, participants were required to write down their thirty most desired positive messages that they would like to receive on social media from their close others (including family members and friends), with the length of 10 to 20 words for each message.

Subsequently, participants were required to vividly imagine reading these messages for one minute per comment and, for each event, to describe their feelings in two sentences. In addition to this, participants were asked to rate the experiences of pleasantness (1-very unpleasant, 9-very pleasant), arousal (1-not aroused at all, 9-very aroused), and how vividly they could imagine receiving these messages (1-very vague, 9-very vivid) on 9-point Likert scales. Twenty messages that could be vividly imagined and induced high levels of pleasantness were selected as positive stimuli for the fMRI experiment. Moreover, experimenters generated 10 neutral messages to be used as messages from a bot (e.g., ‘Maintaining the website environment is everyone’s obligation’), with a length limit of 30 words. For these general messages, each participant applied a 9-point Likert scale to rate the likelihood that such a comment comes from a bot (1-Cannot be from a bot, 9-Definitely from a bot). Within each participant, five general messages that were considered most likely from a bot were applied as neutral stimuli. Finally, combining the interface screenshots and twenty-five messages (20 positive and 5 neutral messages), personalized stimuli were created for each participant for the fMRI experiment to simulate receiving social media messages in close-to-real life. During the fMRI experiment, the individualized messages and interface were displayed. The contents of the messages were identical to those written by each participant previously. In each trial of this task, participants were first presented with the individualized screenshot of the homepage of their social media platform (e.g., Weibo) for 4 seconds, which contained a red bubble floating on the bottom right menu bar, indicating new messages received. There would be a blank (without a red bubble) at the bottom right menu when no new messages were coming in. A jittered black central cross was then displayed on the screen (4-6s). Following this, participants pressed the left button of the response box to jump to the next interface and read the content of the respective message shown in dialogue boxes, once a red central cross appeared on the screen. Dialogue boxes were displayed for 15 seconds, during which participants were required to read and immerse themselves in the emotional experience of receiving and reading these messages (for a previous evaluation of this design, see ref. 47). For those trials without red bubbles (i.e. without new messages received) at the beginning, participants saw the subsequent interface that only contains a blank dialogue box for a jittered duration (4000ms-8000ms) after pressing the button. After each trial, participants rated the pleasantness of the experience (1-very unpleasant to 9-very pleasant) through an interactive rating slider. This was followed by a jittered inter-trial interval (4000ms-6000ms) before the onset of the next trial. Twenty-five messages (20 positive and 5 neutral) assigned to 25 trials were shown in a single run with randomized orders.

For study 3 (taste reward via sweet liquids; Fig. 2c), the procedures were in line with our previously validated experimental procedures that aimed at determining the neural basis of tasting disgusting liquids during fMRI^40^. In the present experiment, participants were administered individualized neutral liquids and sweet liquids to induce a subjective pleasant experience. We employed sweet palatable milk beverages without caffeine, cocoa, fruits, and nuts, to provide a smooth mouthfeel which has been previously shown to engage the striatal reward centers^92^. We provided participants with a selection of five commercially available drinks from evaluated and approved brands to allow a high and individualized pleasantness experience. One day before the experiment, participants tasted 2 ml of each drink and rated how pleasant they felt when tasting them on a 7-point Likert scale (1-not pleasant at all, 7-very pleasant). Two successive tastes were separated by rinsing the mouth with 10ml of water. For each participant, the beverage with the highest rating of pleasantness was used in the following fMRI experiment. Participants who rated all beverages below 5 were not invited to the subsequent experiment. Artificial saliva, rather than clear water, serves as a neutral control because it is colorless and tasteless^93,94^. We prepared artificial saliva of three different concentrations (25mM KCl and 2.5mM NaHCO3; diluted with 30%, 50%, and 80% water respectively) to allow for individualizing the neutral liquid. Each participant was required to taste 10ml of each solution and rate taste intensity (1-no taste at all to 9-very strong taste). Within each participant, the solution with the lowest rating was used later. In total, participants tasted 2 mL of sweet liquid or artificial saliva twenty times (summing up to 40 mL of liquid intake).

During the fMRI experiment, participants lay supine and received visual instructions (Fig. 2c). At the same time, an experimenter stood beside the MRI system and delivered liquids with a pipette, upon receiving auditory instructions from earphones. In each trial, participants first saw a picture of red or blue water drops for two seconds as a visual cue for the upcoming sweet or neutral liquid. This was followed by a jittered central black cross on the screen (3000 ms - 5000 ms). Then, the black cross turned red to inform participants that liquid was being delivered, while the experimenter simultaneously received an auditory instruction to deliver 2 mL of sweet or neutral liquid onto participants’ tongues through a pipette. Participants were asked to press the left button on an MRI-compatible response box when they felt liquid on their tongue for timing purposes, and a ‘taste’ instruction was presented immediately on the screen for 8 seconds, during which participants were required to hold and taste the flavor of the liquid in their mouth and focus on the pleasant or neutral feeling resulting from the liquid. After this, a ‘swallow’ instruction of 5 seconds appeared on the screen to inform participants to swallow the liquid. Then, an auditory instruction asked the experimenter to deliver 10 ml of water to participants to rinse their mouth. This was followed by the ‘rinse’ instruction of 8 seconds and a subsequent ‘swallow’ instruction of 5 seconds visually presented to participants. At the end of a trial, a jittered central black cross (4000 ms - 6000 ms) was displayed as the inter-trial interval. With five trials per run, the tasting experiment had four runs. In line with our previous study^40^, the order of four runs was counterbalanced between consecutive participants as sweet-sweet-neutral-neutral and neutral-neutral-sweet-sweet. At the end of each run, participants rated levels of pleasantness when tasting sweet/neutral liquid in the current run, on a visually presented interactive rating slider that could be manipulated by buttons on the response box.

### Acquisition of MRI data

For studies 1-3, MRI data were acquired on a 3 Tesla MRI scanner (GE MR750, General Electric Medical System). Structural images were obtained through a T1 spoiled gradient recall sequence (scanning parameters: repetition time = 8ms, echo time = 3ms, flip angle=8°, field of view = 256×256mm, voxel size=1×1×1mm, acquisition matrix = 256×256, 176 slices, slice thickness = 1mm) to improve spatial normalization of functional images and exclude participants with apparent brain anomalies. Functional volumes were acquired by using a T2-weighted echo planar imaging (EPI) sequence (scanning parameters: repetition time = 2000ms, echo time = 30ms, flip angle = 90°, field of view = 240×240mm, voxel size = 3×3×3mm, resolution = 64×64, number of slices = 39, slice thickness = 3mm).

### fMRI data pre-processing

Pre-processing for MRI data was implemented with Statistical Parametric Mapping (SPM12, https://www.fil.ion.ucl.ac.uk/spm/software/spm12/). The first five volumes of each run were excluded due to the equilibration of the magnetic field. After that, preprocessing steps for functional images included slice timing correction, realignment for head motion, unwarping to correct magnetic field heterogeneity, co-registration with the skull-stripped and bias-corrected structural image, normalization to the standard Montreal Neurological Institute space (interpolated to 2×2×2 mm voxel size), and spatial smoothing with an 8-mm full width at half maximum Gaussian kernel.

### Behavioral data analysis

Statistical analyses for behavioral data were conducted with JASP software (version 0.19.2; https://jasp-stats.org/). We examined whether participants reported higher levels of experienced romantic love and/or pleasantness in rewarding conditions compared with neutral conditions using two-tailed independent-sample t-tests (See results for details).

### First-level fMRI analysis

A univariate general linear model (GLM) was used to explore the localization of regions involved in the reward processes and to compute beta maps for the decoding models. Participant-level GLMs included task-related regressors modelling brain activity of separate conditions (e.g., study 1: recalling romantic experiences with the current partner, pleasant experiences with a friend, and neutral experiences from time spent by oneself; study 2: reading positive and neutral messages; study 3: tasting sweet and neutral liquids) and one boxcar regressor modelling the rating period to rule out effects of motion. Task regressors were convolved with the canonical hemodynamic response function (HRF), and a high-pass filter of 128 s was employed. Multiple runs of the fMRI task were concatenated for each participant. Outliers of image intensity were detected with CanlabCore tools (https://github.com/canlab/CanlabCore) and were included in the first-level model as additional nuisance variables. Therefore, nuisance variables included: (1) ‘dummy’ regressors representing each run; (2) 24 head motion parameters including the six estimated head movement parameters based on rigid body, their squares, their derivatives and squared derivatives; (3) variables indicating image intensity outliers.

### Multivariate pattern analyses (MVPA)

In line with previous studies^38,40^, we used smoothed data for MVPA as previous studies suggest that smoothing (i.e., 8-mm) can improve inter-subject functional alignment while retaining the sensitivity to mesoscopic activity patterns consistent across participants^95^. To develop multivariate pattern decoders for each study, linear support vector machines (SVMs, *C* = 1) were employed on the averaged whole-brain beta-maps of each participant to decode MRI task conditions (e.g., romantic love versus neutral conditions in study 1). The performance of classification was evaluated by a 10 × 10-fold cross-validation procedure. During this procedure, participants were randomly split into 10 subgroups of equal sample size with the cvpartition function in MATLAB, and each subgroup served as the test data set one-by-one while the remaining nine subgroups were used as the training data set, which was then repeated 10 times to avoid potential bias due to random data splits. The training set and test set were linearly scaled to [-1,1] before entering SVM models. We computed the accuracy of decoding models from receiver operating characteristic curves with two-alternative forced-choice classification, where model response values were compared for brain images of two conditions tested within the same participant (the brain image with a higher model response value was labeled as rewarding, while the other one was labeled as neutral), allowing a static threshold-free classification. Two-sided binomial tests were used to determine whether accuracy was above chance level (50%). Standard errors of accuracy and effect size (Cohen’s *d*) were also reported.

### Identifying brain regions predictive of different types of rewards

To identify core regions for different types of rewarding experiences, we first determined brain regions making reliable contributions to the prediction of decoding models (i.e., model weight maps, also known as backward models). We generated 10,000 bootstrap samples with replacement for each study and performed decoding, which created a null distribution of the weight value for each voxel. The z-scores at each voxel were obtained based on the mean and standard error of this null distribution. The weight maps were then thresholded at multiple thresholds (i.e., uncorrected *P* < 0.05 and *P* < 0.01, as well as FDR *q* < 0.05; two-tailed) on a voxel-wise basis to visualize the extent of clusters while transparently indicating the thresholding level. Secondly, we computed model encoding maps (‘structure coefficients’, also known as forward models) by transforming patterns of decoding models into structure coefficients that identify voxels where the prediction correlated with fMRI activation, making model patterns interpretable^50^. Specifically, one-sample t*-*tests thresholded at multiple thresholds (i.e., uncorrected *P* < 0.05 and *P* < 0.01, as well as FDR *q* < 0.05; two-tailed) were used within each model encoding map to identify voxels associated with each decoding model’s output. Finally, core regions for different types of rewarding experiences were defined as the conjunction of the weight map and the model encoding map, in which voxels reliably contribute to model prediction (that is, model weights) and are related to fMRI activation.

### Comparing neural representations of different rewards

To test whether social and non-social rewards share a neurofunctional basis, we applied a cross-decoding procedure in which each decoding model was used to decode other reward types^56^. A similar neural pattern would allow decoding across the tested models. For these comparisons, monetary reward (study 4, *n* = 39) was also included to determine overlap with secondary non-social rewards. To specifically examine the “common neural currency” hypothesis of reward valuation and to account for heterogeneity in reward modalities and reward targets, we used an ROI-based cross-decoding approach^56^, which focused on core regions of the reward and valuation systems including the striatum and medial frontal cortex. Based on our hypotheses and previous studies^96–98^, we created a mask that encompassed regions of interest (ROIs) involved in reward processes across modalities and reward types^43–45^, including the ventral striatum (VS), dorsal striatum (DS) and medial prefrontal cortex (mPFC), which enables standardized comparisons across reward types. Consistent with prior research^99,100^, the atlas of AAL3^101^ was used to create and fuse masks of VS, DS and mPFC. Specifically, the mPFC mask consisted of functionally validated subregions of mPFC, including the middle cingulate cortex (MCC), anterior cingulate cortex (ACC), medial part of superior frontal gyrus (SFGmedial) and medial orbital part of superior frontal gyrus (PFCventmed), medial orbitofrontal cortex (mOFC) as well as the supplementary motor area (SMA)^102^. We trained specific decoding models on this ROI mask using a 10 × 10-fold cross-validation procedure, yielding accuracy, effect size, and associated *P*-value for model evaluation. Bonferroni correction was applied to adjust for multiple comparisons across a series of cross-decoding accuracy tests against chance level (50%).

Furthermore, river plots were generated to visualize spatial similarity between encoding maps of whole-brain decoding models and ROIs within the reward network. Spatial similarity was computed as cosine similarity between the mask encompassing ROIs and population-level encoding maps within the mask (thresholded at FDR *q* < 0.05 and retaining positive values only for interpretation purposes). To test the selectivity of each ROI to different reward types, relative percentages of each ROI encoded by models were derived.

### Valence specificity, salience and arousal test

To test whether the reward models decode positive emotional experience or simply higher arousal or salience, we tested whole-brain decoding models from studies 1-3 on their responses to arousing positive and negative (versus neutral) movie clips of around 25-second duration in two independent studies (study 5, *n* = 60; study 6, *n* = 36). Pattern response was estimated for each test condition by computing the dot product of each SVM-derived unthresholded model pattern with one participant’s brain activation map, yielding a single scalar value. We tested the sensitivity and specificity of each reward decoding model by examining whether the pattern response to positive movie clips was significantly above zero while the pattern response to negative movie clips was nonsignificant or below zero with one-sample t-tests (one-tailed).

### Generalization of the love decoder to oxytocinergic modulation of love and substance addiction

Previous work has suggested that romantic love is modulated by the neuropeptide oxytocin^1,2^, and a previous study demonstrated that a single dosage of intranasal oxytocin in men enhances subjective pleasantness and the brain’s reward responses to seeing the picture of their romantic female partner^14^. To assess whether the love decoder was sensitive to pharmacologically enhanced experiences of romantic love, we applied the love decoder (restricted to reward system ROIs) to discriminate neural responses to images of participants’ romantic partners from those elicited by unfamiliar opposite-sex individuals under oxytocin and placebo conditions (study 7, *n* = 40). In addition, previous studies have revealed a behavioral and neurobiological similarity between romantic love and addiction, including symptoms of craving, dependence, withdrawal and relapse^5,6,58^. Here, we tested whether substance addiction has shared neural bases with romantic love by applying the decoding model of romantic love restricted to reward system ROIs to distinguish between brain maps of drug users (cannabis users, *n* = 38) and non-users (*n* = 44) when exposed to drug cues (study 8). We also included decoding models of other rewards in this classification analysis to test the specificity of the similarity between love and drug-cue reactivity in contrast to other rewarding experiences.

## Data availability

For studies 1-3, brain patterns and metadata for figures will be shared at https://github.com/Yu-WU-Wyatt/2025-Social-attachment upon publication. The data from study 4 are open-access from a previous study^55^ and are available via https://neurosynth.org/analyses/terms/monetary%20reward/. The data from studies 5-8 were provided by the authors of previous studies^14,36,38^.

## Code availability

Code for reproducing the findings presented in this manuscript is available at https://github.com/Yu-WU-Wyatt/2025-Social-attachment. CanlabCore Tools are available at https://github.com/canlab/CanlabCore.

## Acknowledgements

This work was partly funded by the University Grants Committee Hong Kong (Research Grants Council, HSSPF UGC 37000626; GRF UGC 17617526; GRF UGC 17615525), the Medical Research Fund Hong Kong (HMRF, 23243971), the URC Research Committee (616000) and seed funds of The University of Hong Kong.

## CRediT authorship contribution statement

YW: Conceptualization, Methodology, Formal analysis, Investigation, Visualization, Writing – original draft. XG: Conceptualization, Methodology, Investigation, Data curation, Writing – review & editing. GJ: Conceptualization, Methodology, Investigation, Data curation, Writing – review & editing. RH: Resources, Methodology, Writing – review & editing. DS: Resources, Methodology, Writing – review & editing. XZ: Resources, Data curation, Writing – review & editing. RZ: Resources, Data curation, Writing – review & editing. FZ: Software, Methodology, Writing – review & editing. HJ: Software, Visualization, Writing – review & editing. KF: Investigation, Data curation, Writing – review & editing. JW: Investigation, Data curation, Writing – review & editing. YT: Conceptualization, Writing – review & editing. BB: Conceptualization, Methodology, Supervision, Project administration, Funding acquisition, Writing – review & editing. All authors contributed to the article and approved the submitted version.

---

**Extended Data Fig. 1.**
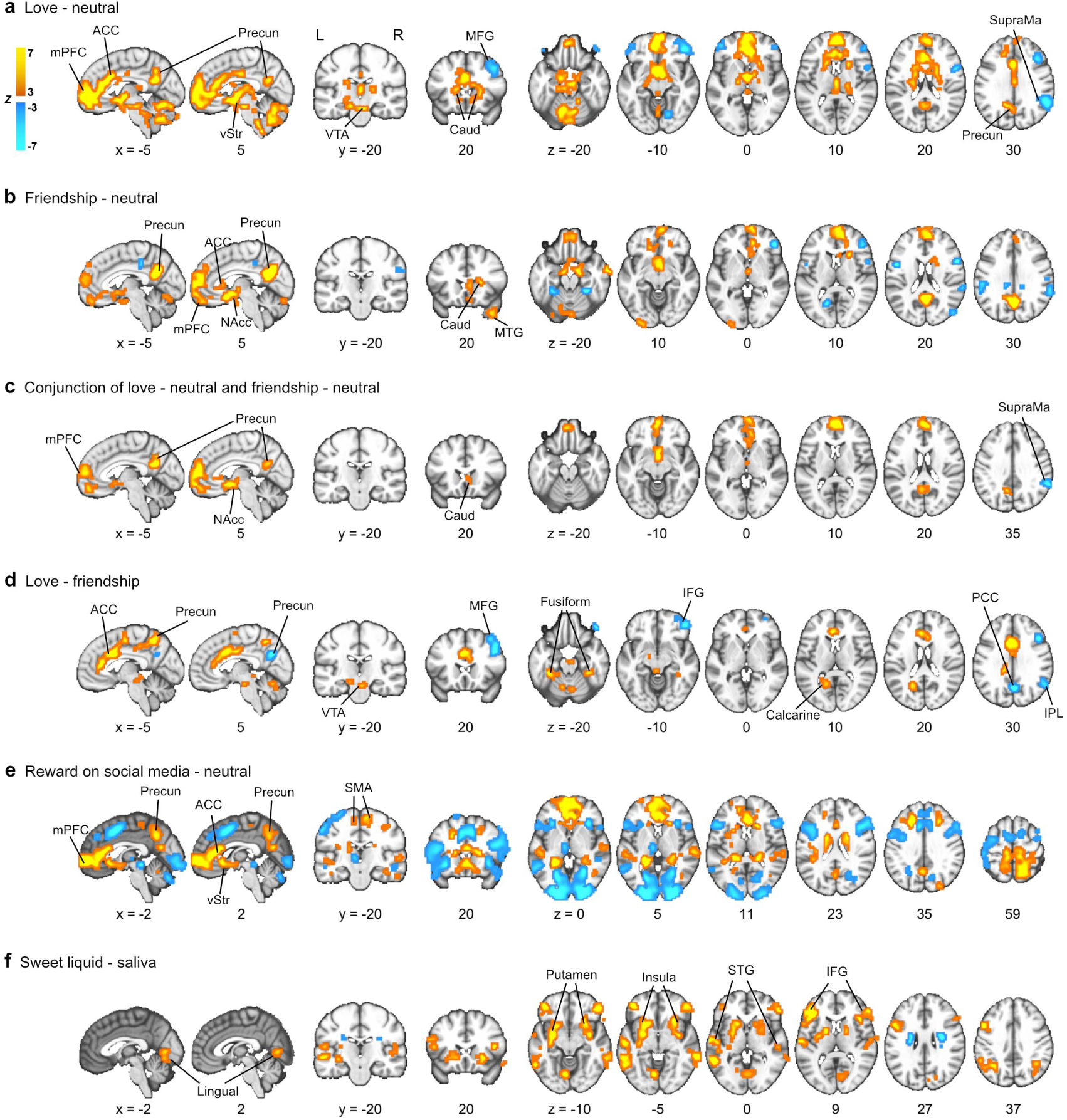
Univariate results for studies 1-3. Activation maps were thresholded at uncorrected *P* < 0.001 for voxels and FDR-corrected *P* < 0.05 for clusters. **a,b,c,d,e,f,** Striatum and frontal cortex, especially the medial prefrontal cortex, are the most consistently engaged brain regions across different reward types. mPFC, medial prefrontal cortex; ACC, anterior cingulate cortex; Precun, precuneus; vStr, ventral striatum; NAcc, nucleus accumbens; VTA, ventral tegmental area; Caud, caudate nucleus; MCC, middle cingulate cortex; STG, superior temporal gyrus; SMA, supplementary motor area; MTG, middle temporal gyrus; IFG, inferior frontal gyrus; MFG, middle frontal gyrus; PCC, posterior cingulate cortex; SupraMa, supramarginal gyrus; L, left hemisphere; R, right hemisphere.

**Extended Data Fig. 2.**
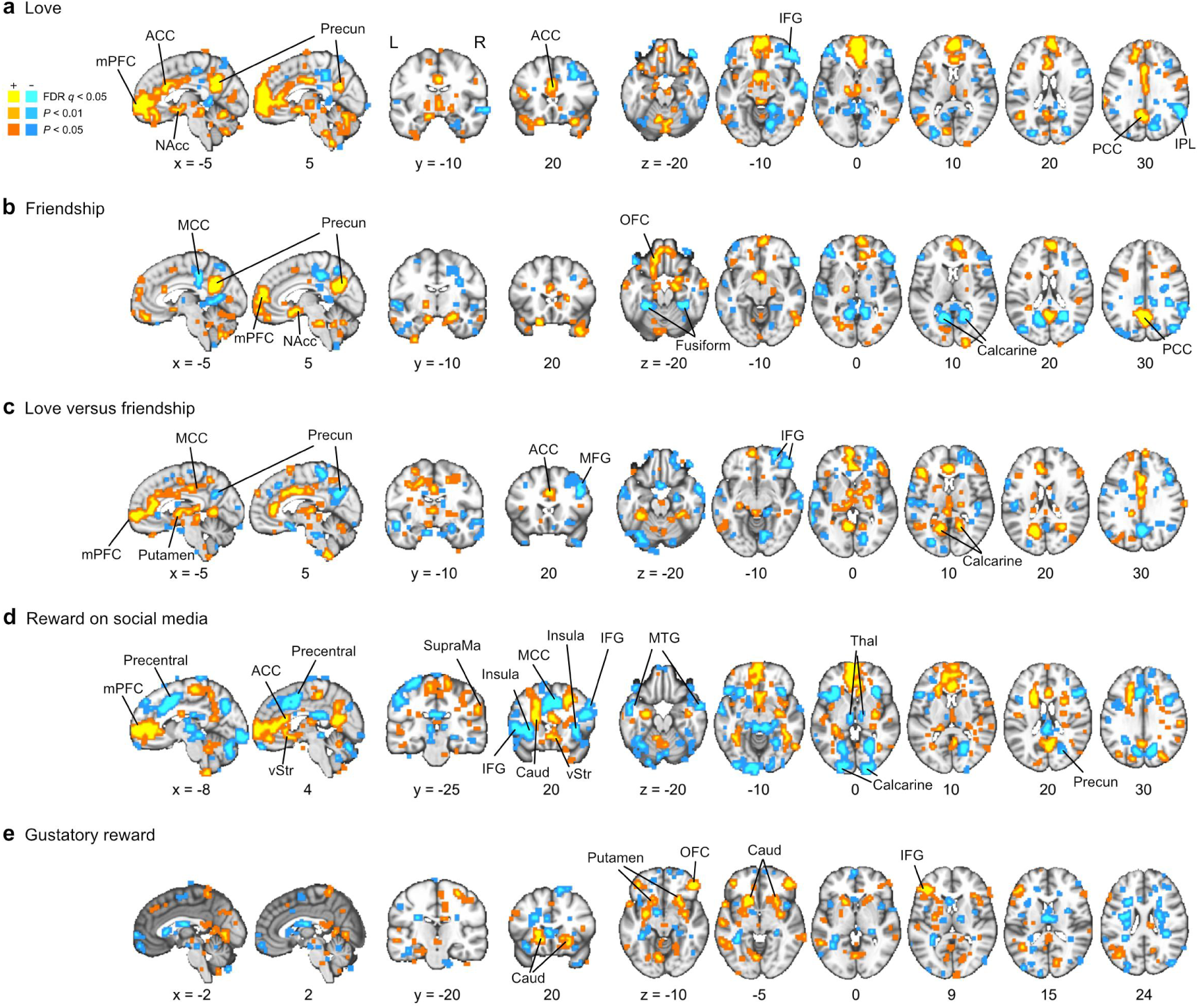
Whole-brain weight maps of decoding models. **a,b,c,d,e,** Whole-brain weight maps of different reward types were thresholded at uncorrected *P*< 0.05 and *P* < 0.01, as well as FDR *q* < 0.05 (two-tailed) to show the extent of clusters. mPFC, medial prefrontal cortex; ACC, anterior cingulate cortex; Precun, precuneus; vStr, ventral striatum; NAcc, nucleus accumbens; OFC, orbitofrontal cortex; MCC, middle cingulate cortex; PCC, posterior cingulate cortex; SupraMa, supramarginal gyrus; Caud, caudate nucleus; MTG, middle temporal gyrus; IFG, inferior frontal gyrus; Thal, thalamus; IPL, inferior parietal lobe; MFG, middle frontal gyrus; L, left hemisphere; R, right hemisphere.

**Extended Data Fig. 3.**
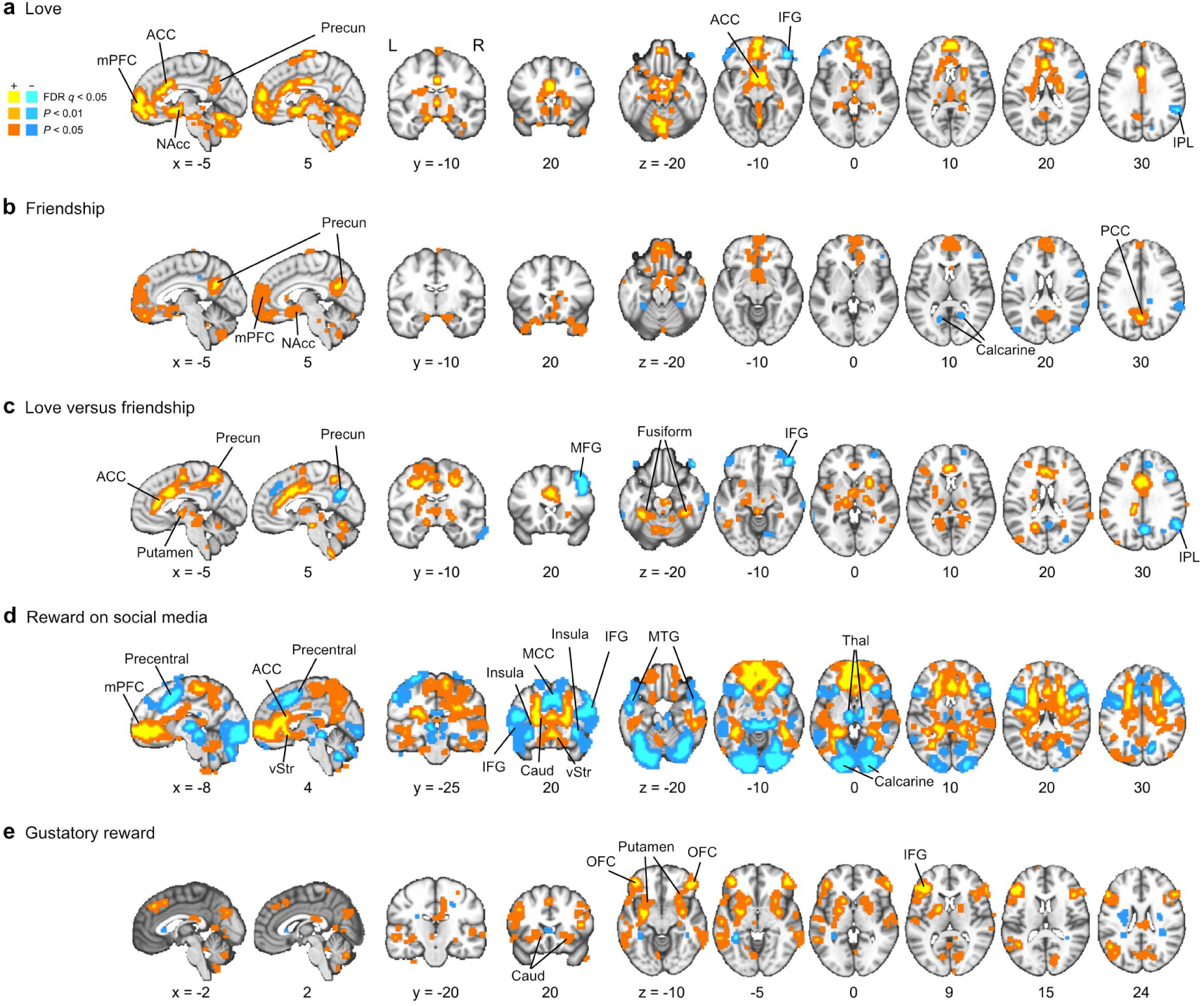
Whole-brain encoding maps of decoding models. **a,b,c,d,e,** Whole-brain encoding maps of different reward types were thresholded at uncorrected *P* < 0.05 and *P* < 0.01, as well as FDR *q* < 0.05 (two-tailed) to show the extent of clusters. mPFC, medial prefrontal cortex; ACC, anterior cingulate cortex; MCC, middle cingulate cortex; Precun, precuneus; vStr, ventral striatum; NAcc, nucleus accumbens; OFC, orbitofrontal cortex; SMA, supplementary motor area; Caud, caudate nucleus; IFG, inferior frontal gyrus; Thal, thalamus. IPL, inferior parietal lobe; MTG, middle temporal gyrus; MFG, middle frontal gyrus; PCC, posterior cingulate cortex; L, left hemisphere; R, right hemisphere.

**Extended Data Fig. 4.**
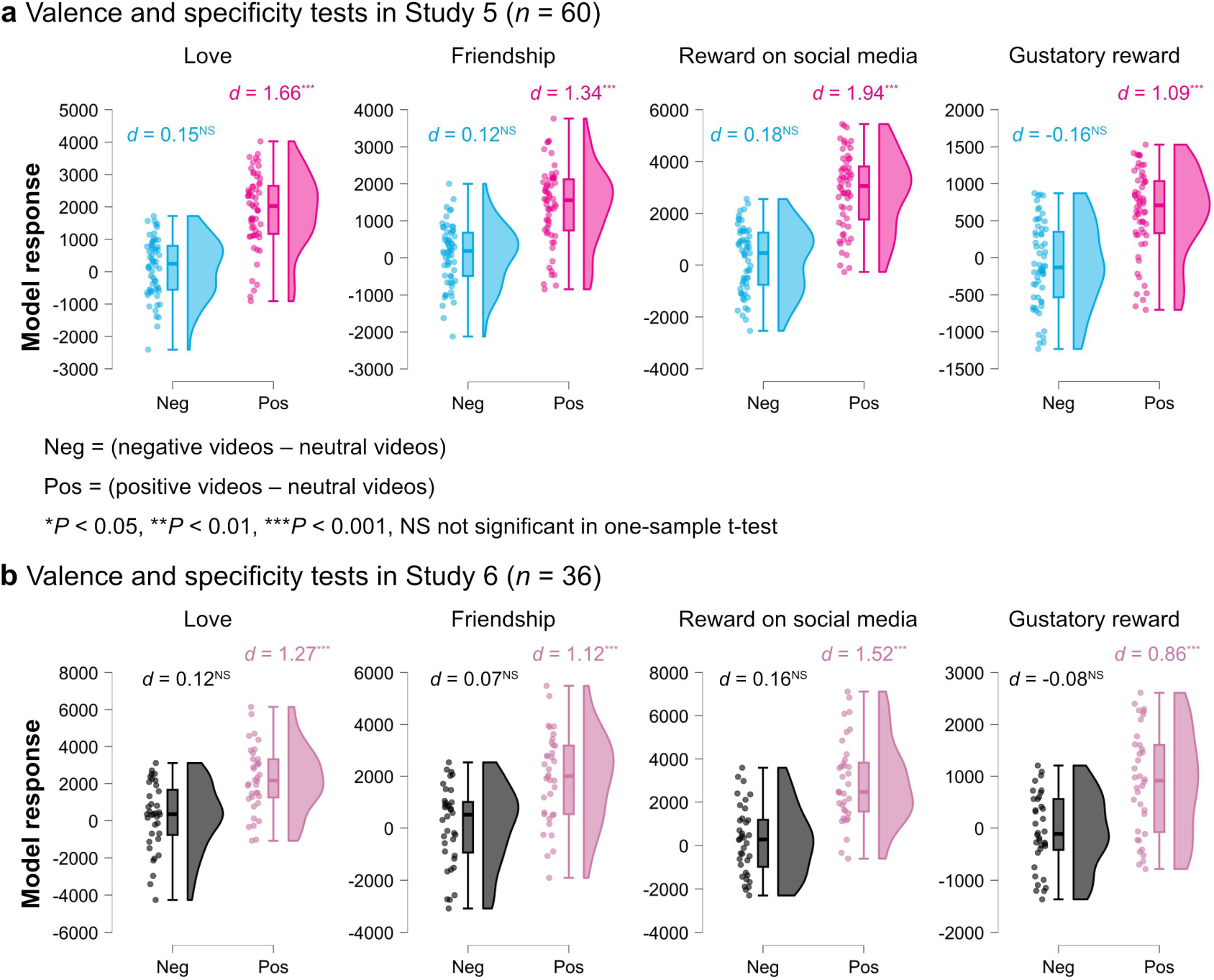
Valence specificity tests in two additional datasets. **a,b,** All whole-brain decoding models showed significant positive responses to highly arousing positive stimuli (Cohen’s *d* > 0.86) but insignificant responses to highly arousing negative stimuli (Cohen’s *d* < 0.18), indicating specificity for positively valenced stimuli. Model responses in studies 5 and 6 were obtained by computing the dot product of each unthresholded weight map of whole-brain decoding models with the brain activation map of each test condition for each participant. Model responses were tested with one-sample t-tests (one-tailed). *d*, Cohen’s *d* effect size; \**P* < 0.05, \*\**P* < 0.01, \*\*\**P* < 0.001, NS not significant. The boundaries of box plots stand for the first and third quartiles, while the whiskers shown are the range of the median ± 1.5 times the interquartile range.

