## Supplementary information for "A neurofunctional signature of romantic love predicts rewards on social media, effects of oxytocin and drug-cue reactivity" (from another version of this preprint: version 2, 2026-08-16)

This file includes:
Supplementary Figures 1-3
Supplementary Table 1
Supplementary Methods
Supplementary References

### Supplementary Figures

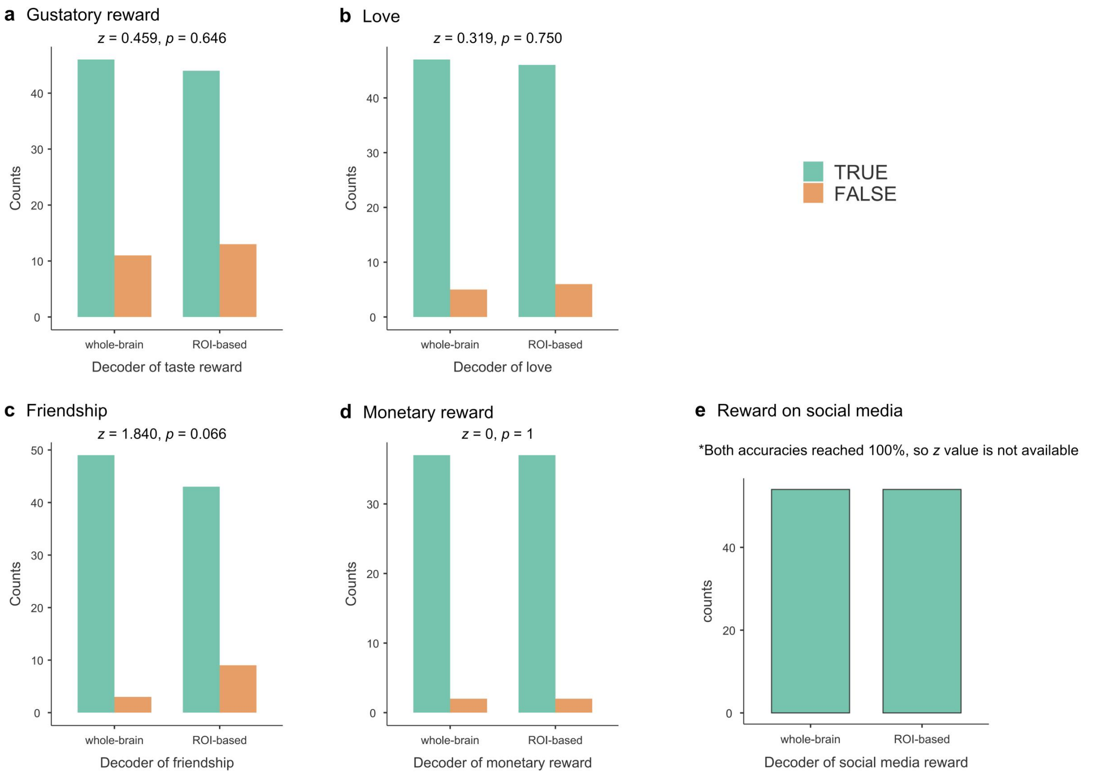


**Supplementary Figure 1. Accuracy differences between whole-brain and ROI-based decoding models. a,b,c,d,e,** Bar plots show counts of true or false outputs of whole-brain and ROI-based decoding models. *Z* tests for difference in two proportions are used to test accuracy differences between whole-brain and ROI-based decoding models. Accuracy of both whole-brain and ROI-based decoding models for reward on social media reached 100%, so the *Z* value is not applicable. Accuracy levels of the whole-brain and the ROI-based decoding models for each type of reward were similar. However, we observed a pronounced reduction in decoding accuracy for decoders of friendship when restricting the decoder to ROIs relative to its whole-brain counterpart, although this difference did not reach statistical significance ($p = 0.066$). This pattern suggests that neural signatures of friendship are distributed beyond the reward network and may also involve regions such as the precuneus and posterior cingulate cortex.

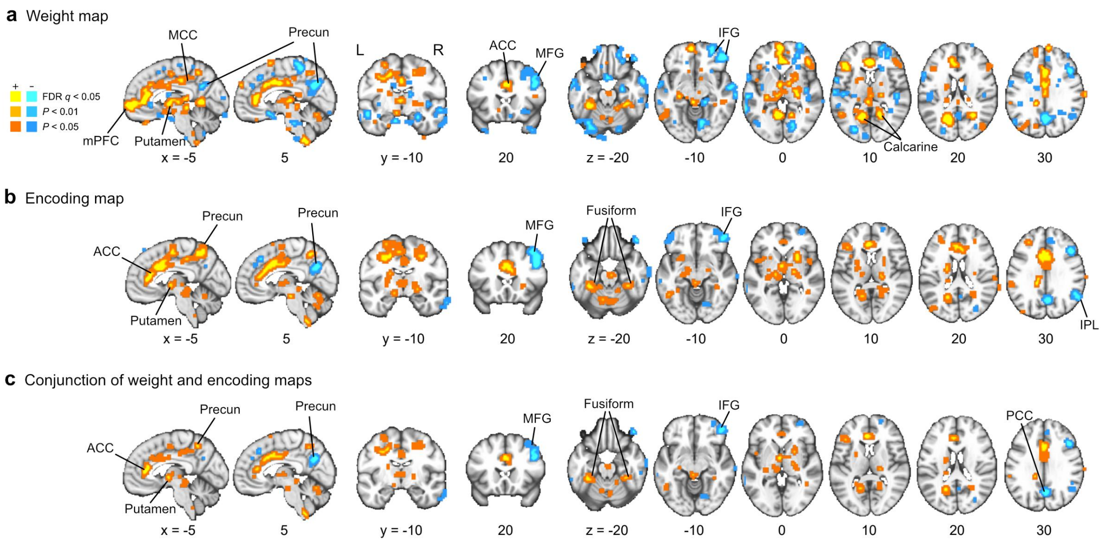


**Supplementary Figure 2. Core brain systems distinguishing love from friendship with pleasantness level as the covariate.** Given that participants reported higher pleasantness levels in the love condition compared with the friendship condition, we ran additional analyses that included pleasantness level as the covariate. Results showed that the decoding accuracy (love versus friendship) remained robust after accounting for pleasantness level (accuracy = 83% ± 5.2% SE, $P < 10^{-5}$, Cohen's $d = 1.54$). Furthermore, the core brain systems distinguishing love from friendship were also consistent with the findings without taking pleasantness level as the covariate (See Supplementary Fig. 2c and Fig. 5c). The whole-brain decoding model of love versus friendship here was developed through the identical procedure of 10 × 10-fold cross-validation specified in the main text. **c,** Core brain systems were defined as the conjunction of **a,** the weight map, and **b,** the model encoding map. These maps were thresholded at uncorrected $P < 0.05$ and $P < 0.01$, as well as FDR $q < 0.05$ (two-tailed) to show the extent of clusters.

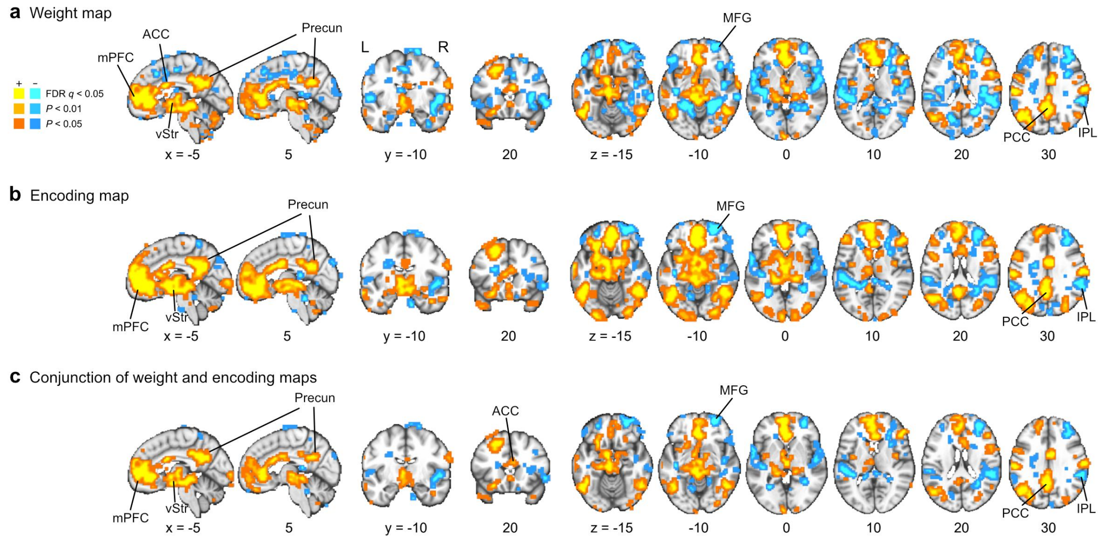


**Supplementary Figure 3. Core brain systems of drug-cue reactivity.** The whole-brain decoding model of drug-cue reactivity was developed through the identical procedure of 10 × 10-fold cross-validation specified in the main text, with a neutral condition of viewing neutral images. **c,** Core brain systems were defined as the conjunction of **a,** the weight map, and **b,** the model encoding map. These maps were thresholded at uncorrected *P* < 0.05 and *P* < 0.01, as well as FDR *q* < 0.05 (two-tailed) to show the extent of clusters. Univariate results of drug-cue reactivity can be found in Supplementary ref. 4.

### Supplementary Tables

Supplementary Table 1. Information for each study.

| Study | Dataset ref. | Stimuli Types | *N* (female) | Mean age | Study location | MRI system |
|---|---|---|---|---|---|---|
| 1 | This paper[a] | Visual cues of keywords for recalling past experiences | 52(28) | 20.16 | China | 3T GE |
| 2 | This paper[a] | Images of social media interface | 54(30) | 20.73 | China | 3T GE |
| 3 | This paper[a] | Sweet liquids and artificial saliva | 57(33) | 19.82 | China | 3T GE |
| 4 | Speer et al., 2023[b,1] | Images of monetary reward | 39(28) | 23.62 | Finland | 3T Siemens Verio |
| 5 | Zhang et al., 2025[c,2] | Positive and negative videos | 60(23) | 19.55 | China | 3T GE |
| 6 | Zhang et al., 2025[c,2] | Positive and negative videos | 36(17) | 21.03 | China | 3T GE |
| 7 | Scheele et al., 2013[d,3] | Face images of one's romantic partner and unknown others from the opposite sex | 40(0) | 26.03 | Germany | 3T Siemens Trio |
| 8 | Zhou et al., 2019[b,4] | Images of cannabis | 82(0)* | 23.02 | Germany | 3T Siemens Trio |

[a]Current study data; [b]The brain maps were obtained at https://osf.io/ydfrs/; [c]Data from our lab; [d]We applied for the sharing of anonymized MRI data from the authors. *Participants consisted of 38 drug users and 44 healthy controls. Participants from all other studies were healthy adults.

### Supplementary Methods

#### Questionnaires

Two questionnaires were applied to evaluate participants' eligibility for Study 1:

1) Passionate Love Scale[5]. This scale measures the intensity of passionate love. There are 30 items (e.g., "I can feel my tenderness for him (her).") to be graded in 9 response anchors from 1 (Not at all true) to 9 (Definitely true).

2) McGill Friendship Questionnaire[6]. This questionnaire comprises two parts: McGill Friendship Questionnaire-Respondent's Affection (MFQ-RA) that assesses friendship satisfaction with 16 items and McGill Friendship Questionnaire-Friend's Function (MFQ-FF) evaluating six friendship functions with 30 items (that is, stimulating companionship, help, intimacy, reliable alliance, self-validation, and emotional security). The respondent indicates the degree of agreement for items on a 9-point scale (from -4 "Very Much Disagree" to +4 "Very Much Agree") for MFQ-RA, while reporting the frequencies of behaviors stated in items on an 8-point scale (from 0-never to 8-always) for MFQ-FF.
